# Structural basis of substrate recognition and allosteric inhibition in human B^0^AT2

**DOI:** 10.64898/2026.06.16.732524

**Authors:** Yunlei Cao, Yunlong Cao, Deqiang Yao, Shaobai Li, Qian Wang, Shaohua Shi, Futang Wan, Mingyue Li, Sijia Huang, Huaisheng Lu, Qin Yang, Mi Cao, Yafeng Shen, Chao Zheng, Shanshuang Chen, Wenqing Xu, Jing Xue, Jian Wu, Pengfei Lan, Ming Lei

**Author notes:** Correspondence (M.Lei.), (P.L.), (J.W.), (J.X.). These authors contributed equally to this work (Yunlei Cao, Yunlong Cao).

## Abstract

The SLC6 family is a major target for neuropsychiatric therapeutics. Human B^0^AT2 (SLC6A15) regulates amino acid homeostasis and glutamatergic transmission and is linked to major depressive disorder, yet its transport and inhibition mechanisms remain unclear. Here we report cryo-EM structures of B^0^AT2 in apo state and in complex with substrates (proline, leucine, methionine) and inhibitors (loratadine, tiagabine), capturing outward-open, early substrate-bound intermediate, outward-occluded, and inward-open conformations along the transport cycle. These structures reveal an early substrate-sensing mechanism at the substrate-binding pocket (S1), where a substrate-dependent rotameric switch of Phe308 remodels pocket geometry to tune substrate accommodation and selectivity. Loratadine stabilizes an outward-occluded state via allosteric inhibition at the extracellular S2 pocket, whereas tiagabine stabilizes the inward-open state through cooperative multi-site inhibition involving S1 and two previously unrecognized intracellular cavities (S3 and S4). Together with functional assays, these data define the molecular basis of B^0^AT2 substrate selectivity and state-dependent inhibition. Notably, the two intracellular cavities are conserved across SLC6 transporters, reflecting a shared intracellular vestibular architecture and enabling rational design of conformation-selective modulators for neuropsychiatric disorders.

## Introduction

The solute carrier 6 (SLC6) family, also known as the neurotransmitter-sodium symporter (NSS) family, comprises a major class of transporters that mediate the uptake of key neurotransmitters and play essential roles in diverse physiological processes, particularly in psychiatric disorders (Kristensen, Andersen et al., 2011, Pramod, Foster et al., 2013). Prominent members include the serotonin transporter (SERT), norepinephrine transporter (NET), dopamine transporter (DAT), GABA transporter 1 (GAT1), and glycine transporter 1 (GlyT1), which are major targets for antidepressants, anticonvulsants, illicit drugs, and antipsychotics (Kristensen et al., 2011, Navratna & Gouaux, 2019). B^0^AT2 (SLC6A15, also known as V7-3) is a member of the SLC6 subfamily and functions as a sodium-dependent neutral amino acid transporter predominantly expressed in the central nervous system, including glutamatergic neurons and astrocytes (G.R. Uhl, 1992, Santarelli, Wagner et al., 2016, Ulrich, Hägglund et al., 2013). Its expression is enriched in brain regions associated with emotional regulation, behavior, and feeding control (Alquier, Drgonova et al., 2013, Chandra, Francis et al., 2017, Santarelli et al., 2016). Functionally, B^0^AT2 preferentially transports methionine, proline, and branched-chain amino acids including leucine, isoleucine, and valine (Bröer, Tietze et al., 2005, Takanaga, Mackenzie et al., 2005). These substrates serve as essential precursors for neurotransmitter synthesis, contribute to the amino acid metabolism in the brain, and are implicated in glutamatergic signaling and related metabolic pathways **(Fig. 1A)** (Santarelli, Namendorf et al., 2015, Santarelli et al., 2016). Genetic studies connect B^0^AT2 to stress susceptibility and mood-related phenotypes (Chandra et al., 2017, Kohli, Lucae et al., 2011, Szopa, Poleszak et al., 2018, Wang, Liu et al., 2017), yet the underlying mechanism remains unclear.

**Figure 1.**
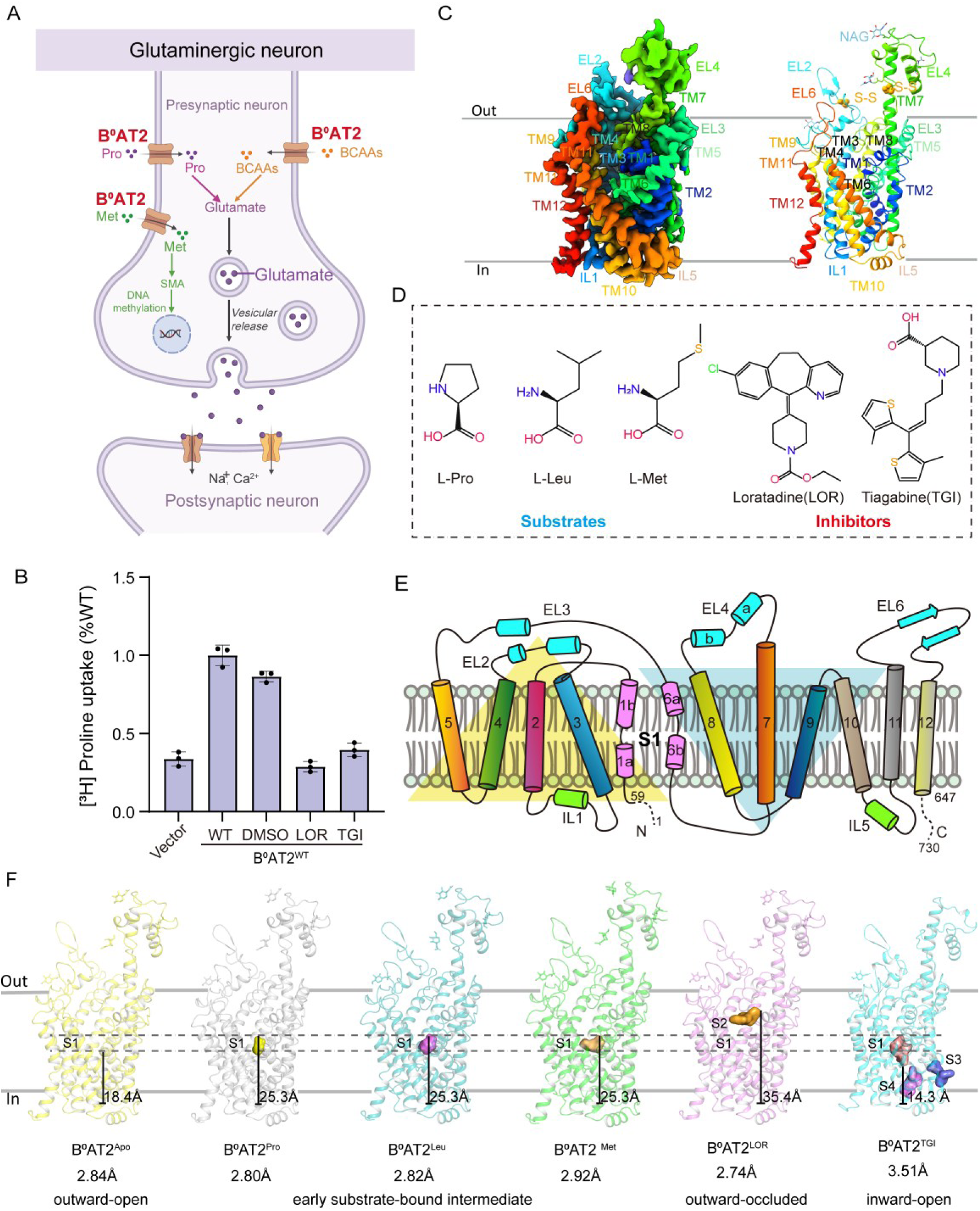
Functional characterization and structure of B^0^AT2. (**A**). Schematic representation of B^0^AT2-mediated transport of neutral amino acids (Pro, Leu, Met) and their metabolic pathways in glutamatergic neurons. (B) Inhibition of [3H]-proline uptake by LOR and TGI in WT B^0^AT2 and vector served as controls. Data were normalized to WT B^0^AT2. Data represent mean ± s.d. (error bars), n = 3 technical replicates. The experiment was performed independently twice with similar results (C) Overall structure of B^0^AT2. N terminus, C terminus, TM1-12, and extracellular or intracellular loops are labeled. (**D**). Chemical structures of selected B^0^AT2 substrates and inhibitors. (E) Topology diagram of B^0^AT2, which adopts a LeuT-fold architecture, from the N-terminal Asp59 to the C-terminal Asp647. Unresolved regions are shown with dashed lines. (F) Overall structures of B^0^AT2 at apo state (B^0^AT2^Apo^) and binding with Pro (B^0^AT2^Pro^), Leu (B^0^AT2^Leu^), Met (B^0^AT2^Met^), LOR (B^0^AT2^LOR^), TGI (B^0^AT2^TGI^) are represented by yellow, blue-white, teal, green, violet, and cyan, respectively. The orthosteric site S1 and the allosteric sites S2, S3 and S4 are indicated, along with their respective distances from the cytoplasmic membrane plane. The corresponding conformational states are annotated below.

As depression and anxiety remain the leading causes of disability worldwide, there is sustained interest in elucidating the molecular mechanisms that couple metabolism to neuronal signaling (Kohli et al., 2011, Wang et al., 2017). Current treatments are effective in only a subset of patients and are limited by delayed onset, interindividual variability, and adverse effects (Cui, Li et al., 2024). In this context, B^0^AT2 is of particular interest because it regulates the availability of neutral amino acids in the brain and has been linked to glutamatergic homeostasis, stress responses, and depression-related phenotypes (Santarelli et al., 2015, Santarelli et al., 2016). Despite this growing biological and pharmacological interest, the structural basis of substrate selectivity, ligand recognition, and conformational coupling in B^0^AT2 has remained unclear in the absence of high-resolution structures (Chandra et al., 2017, Kukulowicz, Pietrzak-Lichwa et al., 2023). Only a small number of non-amino-acid inhibitors have been reported, including LOR, desloratadine analogues, and TGI (Cuboni, Devigny et al., 2014, Kukulowicz, Siwek et al., 2025). LOR was identified as the first characterized B^0^AT2 inhibitor (Cuboni et al., 2014), whereas TGI, a clinically used GAT1 inhibitor, has been shown to inhibit both B^0^AT2 and SLC6A20 (Broer, Hu et al., 2024, Kukulowicz et al., 2025). These compounds serve as chemical probes to elucidate the molecular mechanisms underlying B^0^AT2 inhibition.

In this study, we determined a series of cryo-electron microscopy (cryo-EM) structures of human B^0^AT2 in the apo state (B^0^AT2^Apo^), bound to the substrate proline (B^0^AT2^Pro^), leucine (B^0^AT2^Leu^), and methionine (B^0^AT2^Met^), or in complex with inhibitors LOR (B^0^AT2^LOR^) and TGI (B^0^AT2^TGI^). These structures capture multiple states along the transport cycle, and together with [³H]-proline uptake assays and mutational analyses, define the molecular basis of substrate recognition, conformational tuning of the substrate-binding pocket, and state-dependent inhibition of B^0^AT2. Our structures also reveal two intracellular cavities (S3 and S4) that are preferentially exposed in the inward-open state, uncovering previously unrecognized intracellular regulatory features, and provide structural insights into the conformation-selective modulation of B^0^AT2 and related NSS transporters.

## Results

### Functional characterization and structural architecture of B_0_AT2

To investigate the structural basis of substrate recognition and transport, human B^0^AT2 was expressed in HEK293T cells, and its plasma membrane localization and transport activity were examined by confocal microscopy and [³H]-proline uptake assays **(Fig. 1B; Fig. EV1A)**. After expression construct optimization and detergent screening (**Fig. EV1B–G**), cryo-EM structures of B^0^AT2 were determined in the apo state and in complex with substrates (Pro, Leu, and Met) or inhibitors (LOR and TGI) at resolutions of 2.74–3.51 Å (**Fig. 1C–D, F; Fig. EV1–2; Table. EV1**). These structures of B^0^AT2 reveal the ligand-binding modes and conformational states along the transport cycle.

The B^0^AT2^Apo^ structure adopts a compact architecture, exhibiting the canonical LeuT (leucine transporter)-fold composed of 12 transmembrane (TM) helices with cytoplasm-facing N- and C-termini **(Fig. 1C, E)** (Nyola, Karpowich et al., 2010, Yamashita, Singh et al., 2005). B^0^AT2^Apo^ exhibits a pseudo-twofold symmetry relating TM1–5 to TM6–10, with unwound segments in TM1 and TM6 forming the primary substrate-binding pocket (S1) near the symmetry axis **(Fig. 1E)**, while the C-terminal segment of TM7 extends toward the extracellular side (**Fig. 1C, E**). The extracellular vestibule above the S1 pocket remains solvent-accessible while the intracellular vestibule is closed, indicative of an outward-open conformation (**Fig. 1C**). As in other LeuT-fold transporters, B^0^AT2 comprises a mobile core region (TM1, TM3, TM6, TM8) and a surrounding stabilizing domain, which together mediate the conformational changes during substrate transport (**Fig. 1C, E**) (Penmatsa & Gouaux, 2014). Several extracellular loops (EL) including EL2, EL4, and EL6 form the vestibule above the substrate-binding cavity, while short intracellular loops (IL) such as IL1 and IL5 line the cytoplasmic side of B^0^AT2 (**Fig. 1C**). EL2 and EL4 contain four N-linked glycosylation sites (Asn213, Asn383, Asn394, and Asn424), and are stabilized by two disulfide bonds (Cys182-Cys197 and Cys367-Cys426) (**Fig. 1C**). Sequence comparison across LeuT and representative NSS transporters reveals that these loops are less conserved than the transmembrane core (**Fig. EV3**) (Kukulowicz et al., 2023, Takanaga et al., 2005), suggestive of their diversified roles among SLC6 family transporters. Likely due to the structural flexibility, both the N-terminal (residues 1-58) and C-terminal (residues 648–730) regions are unresolved in the cryo-EM density (**Fig. 1E**). Taken together, the B^0^AT2 apo structure in an outward-open state provides a framework for further understanding substrate recognition and allosteric inhibition of B^0^AT2.

### Structural basis of substrate recognition and selectivity

B^0^AT2 is a Na⁺-dependent transporter that selectively mediates the uptake of a restricted subset of neutral amino acids, most prominently Pro, Met, and branched-chain amino acids (Leu, Ile, and Val), while exhibiting a minimal activity toward charged, aromatic, or monoamine substrates (Bröer et al., 2005, Takanaga et al., 2005). To elucidate the mechanism of substrate selectivity, we determined the cryo-EM structures of human B^0^AT2 in complex with Pro, Leu, and Met. In all three structures, amino acids occupy the S1 substrate-binding pocket located ∼18 Å below the extracellular membrane surface (**Fig. 2A–C; Fig. EV4A, 4I)**. The pocket is primarily formed by residues from TM1, TM3, TM6, and TM8, with the backbone in the unwound regions of TM1 and TM6 coordinating the α-amino and α-carboxyl groups of the substrates (**Fig. 2B–C; Fig. EV4B**). The densities of two putative Na⁺ ions are identified near the junction of TM1a and TM1b, consistent with the sodium sites observed in other LeuT-fold transporter structures (**Fig. 2C; Fig. EV3, EV4A–H**) (Kristensen et al., 2011, Navratna & Gouaux, 2019, Penmatsa & Gouaux, 2014, Pramod et al., 2013, Wei, Li et al., 2024). Na-1 coordinates Asn83 and the substrate carboxyl group (**Fig. 2C; Fig. EV4C**), while Na-2 lies ∼6 Å from the substrate α-carbon and is coordinated by the sidechain of Ser474 and the backbone carbonyls of Gly76, Val79, and Leu470 (**Fig. 2C; Fig. EV4D**). Mutations of residues for Na⁺ coordination (N83A, N341A, and S474A) reduce the transport activity in [³H]-proline uptake assays (**Fig. 2D**), underscoring the importance of the sodium-substrate coupling (Takanaga et al., 2005, Yamashita et al., 2005).

**Figure 2.**
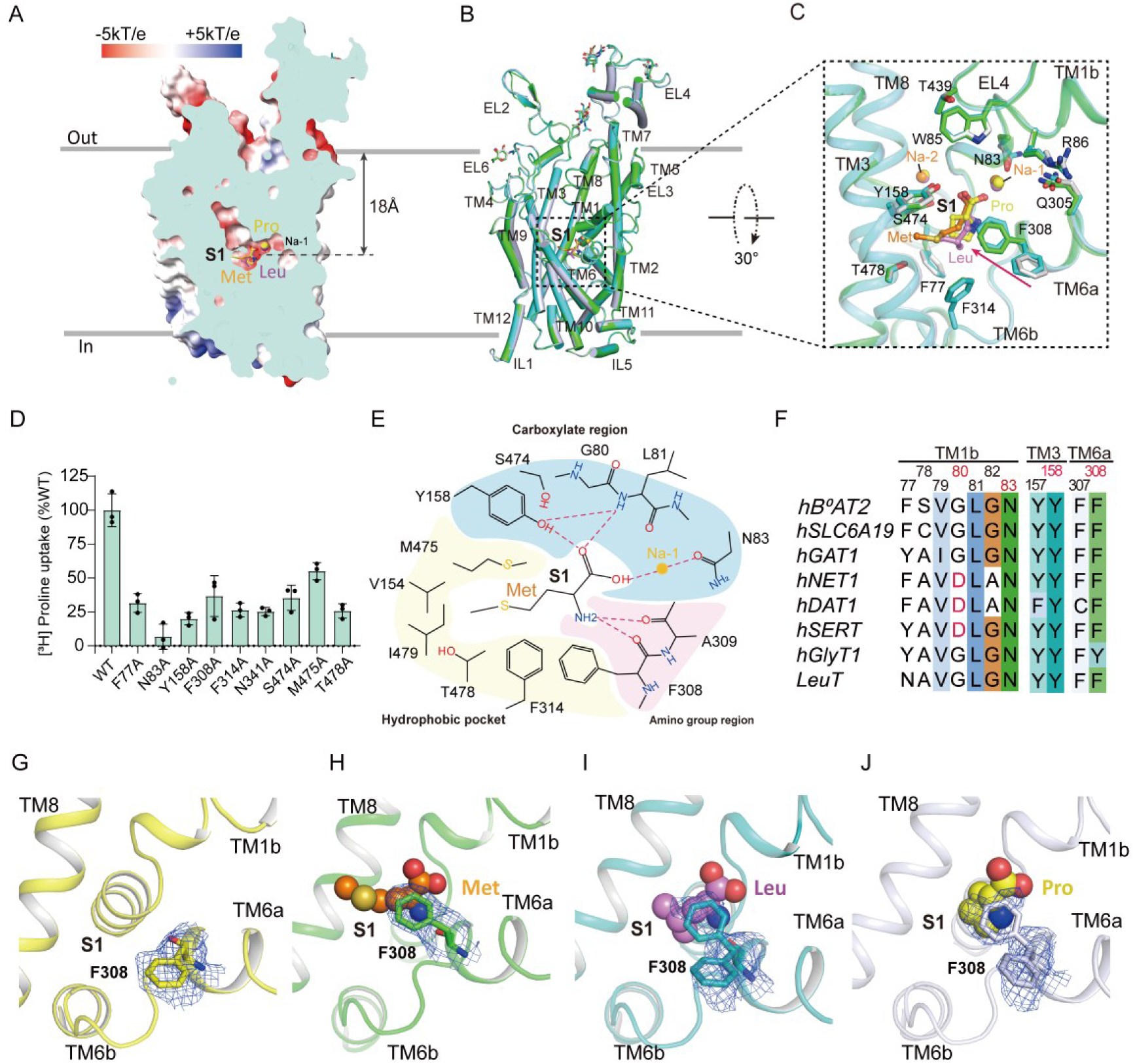
Structural basis of substrate recognition and selectivity through Phe308-mediated tuning of the S1 pocket. (A). Cut-open electrostatic surfaces of B^0^AT2 bound to Pro, Leu and Met, shown as sticks. Electrostatic potential calculated in PyMOL/APBS (red to blue, −5 to +5 kT e⁻¹). Distances from the extracellular membrane indicated. (B). Superposition of B^0^AT2^Pro^, B^0^AT2^Leu^ and B^0^AT2^Met^, shown in cylindrical representation and colored blue-white, teal, and green, respectively; substrates in sticks (yellow, violet, and brown) occupy the canonical S1 pocket. Corresponding TM helices labeled. (C). Close-up view of the substrate-binding pocket. Substrate-interacting residues are shown as sticks and colored to match the corresponding substrate-bound structures, while Pro, Leu and Met are depicted as sticks. Modeled Na⁺ ions are shown as spheres and colored consistently with their associated substrates. (D) Transport activity of wild-type (WT) B^0^AT2 and mutants quantified using [3H]-proline uptake assay. The transport activity of each mutant is normalized to that of WT B^0^AT2. The uptake duration is 1 min to ensure the uptake occurs within the linear range. Data represent mean ± s.d. (error bars), n = 3 technical replicates. The experiment was performed independently twice with similar results. (E) Road-kill plot of Met interactions within the S1 pocket. Hydrogen bonds and coordination interactions are shown as red dashed lines. The amino group region, carboxylate region, and hydrophobic pocket are colored pink, teal, and pale yellow, respectively. Modeled Na-1 is shown as yellow spheres. (F) Sequence alignment of B^0^AT2 with representative NSS transporters, SLC6A19 and LeuT shows that key substrate-binding residues, including Phe308, Tyr158, Gly80, as well as Na-1-coordinating residues such as Asn83, are highly conserved. (**G–J**) Cryo-EM densities of Phe308 (sticks with blue mesh) in different substrate-bound complexes. An alternative Phe308 rotamer was modeled in B^0^AT2^Leu^ and B^0^AT2^Pro^. B^0^AT2^Apo^ (outward-open) is shown in comparison with B^0^AT2^Met^, B^0^AT2^Leu^, and B^0^AT2^Pro^, which represent early substrate-bound intermediate states. All structures are shown in cartoon representation, with substrates displayed as spheres (orange, violet, and yellow).

Consistent with other amino acid-transporting LeuT-fold transporters, the B^0^AT2 S1 pocket can be functionally described as comprising three spatially distinct regions: an amino group region, a carboxylate region, and a hydrophobic pocket, which together mediate substrate binding (**Fig. 2E; Fig. EV4B**) (Wei et al., 2024, Yamashita et al., 2005). At the amino group region of B^0^AT2, the substrate amino groups interact with the backbone carbonyls of Phe308^TM6^ and Ala309^TM6^ (**Fig. 2E; Fig. EV4B**). At the carboxylate region, substrate carboxyl groups are coordinated by Na-1, the backbone amide of Leu81^TM1^, and the hydroxyl group of Tyr158^TM3^ (**Fig. 2E; Fig. EV4B**). The sidechain of Tyr158^TM3^ forms a hydrogen bond with the backbone of Leu81, stabilizing the unwound segment of TM1 (**Fig. 2E; Fig. EV4B**). Notably, substitution of Tyr158 with an alanine residue severely impairs the transport activity of B^0^AT2 (**Fig. 2D**). Consistent with these observations, a tyrosine residue is highly conserved at the equivalent positions among all SLC6 family transporters (**Fig. 2F**) (Kristensen et al., 2011, Li, Wang et al., 2024, Wei et al., 2024). In contrast, a neighboring residue in the carboxylate region of B^0^AT2, Gly80, differs from those of monoamine transporters, which often contain acidic residues at the equivalent sites (such as Asp46 in Drosophila dopamine transporter (dDAT) and Asp98 in SERT) (**Fig. 2F; Fig. EV5A–E**) (Coleman, Green et al., 2016, Wang, Penmatsa et al., 2015). Therefore, we conclude that the relatively less negatively charged environment of B^0^AT2 carboxylate region is suited to neutral amino acid recognition whereas the acidic sidechains in the corresponding regions of monoamine transporters are better for binding of positively charged amine substrates (Kristensen et al., 2011, Pramod et al., 2013, Wei et al., 2024). The side chains of substrates (Met, Leu, and Pro) are accommodated within the hydrophobic region of the S1 pocket, which is formed by a panel of hydrophobic and aromatic residues (**Fig. 2C, E**). Alanine substitutions of these residues markedly reduce the transport activity (**Fig. 2D**), underscoring their importance in substrate side-chain recognition of B^0^AT2. Structural comparison revealed that the overall dimensions of the hydrophobic pocket are similar in all three substrate-bound B^0^AT2 structures (**Fig. 2A–C**), indicating that this region of B^0^AT2 is structurally compatible only with side chains of moderate size with a hydrophobic character, but not compatible with polar, charged, and bulky side chains. Consistent with this idea, structural modeling of a Met-to-Trp substitution generates severe steric clashes within the hydrophobic pocket of B^0^AT2 (**Fig. EV5F–G**). Taken together, these analyses define a structurally and chemically constrained S1 pocket that establishes the basis for substrate selectivity and sets the stage for subsequent conformational tuning during substrate binding.

### Phe308 acts as a key conformational sensor during substrate capture

Structural comparison of apo and substrate-bound B^0^AT2 structures revealed that the overall architecture remains virtually unchanged upon substrate binding, with the only discernible difference being a pronounced rotameric rearrangement of Phe308 that locally remodels the S1 binding pocket **(Fig. 2G–J)**. In the apo state, Phe308 adopts a rotamer positioned away from the substrate pocket, maintaining an open extracellular connection, in alignment with an outward-open conformation (**Fig. 2G**). In contrast, In the B^0^AT2^Met^ structure, the phenol ring of Phe308 rotates to cap the hydrophobic cavity, occluding the S1 pocket, leaving only a narrow gap above the substrate (**Fig. 2H; Fig. EV4I; Movie EV1)**, suggesting a direct role in substrate capture. In both the B^0^AT2^Leu^ and B^0^AT2^Pro^ structures, the cryo-EM densities for Phe308 reveal a dual-conformation, corresponding to the apo-like and B^0^AT2^Met^-like states, respectively (**Fig. 2I–J; Fig. EV4I**), suggesting that both substrate-free and substrate-bound states were co-captured during sample preparation. Importantly, the conformational state of Phe308 is tightly coupled to substrate occupancy, consistent with a direct response to ligand binding. Although all substrate-bound samples were prepared under identical conditions, Phe308 displays distinct conformational distributions. Compared with B^0^AT2^Leu^ and B^0^AT2^Pro^, B^0^AT2^Met^ shows a markedly more defined density for the capping rotamer of Phe308, suggesting preferential stabilization of this conformation by methionine and implying a higher apparent affinity, consistent with previous electrophysiological data(Takanaga et al., 2005). These results indicate that Phe308 sensitively discriminates between substrates and contributes to substrate recognition and capture.

In canonical LeuT-fold transporters, substrate binding triggers large-scale rearrangements of the extracellular gating network, including movements of TM1b and TM6a, as well as local rearrangements of conserved gating residues such as Phe308 and Tyr158, and closure of the Arg86–Asp525 gate pair (**Fig. 2F; Fig. EV6A**), leading to partially occluded or fully occluded states(Cheng & Bahar, 2014, Wang et al., 2015, Wei et al., 2024, Yamashita et al., 2005) (**Fig. EV6B-C**). For example, both the partially-occluded dDAT^DCP^ structure (PDB: 4XPA) and the fully occluded LeuT^Leu^ structure (PDB: 2A65) display such global rearrangements (**Fig. EV6B–C)**(Wang et al., 2015, Yamashita et al., 2005). Consistent with this conserved role, alanine substitution of these gate residues markedly reduces B^0^AT2 transport activity (**Fig. 2D; Fig. EV6D**). However, substrate-bound B^0^AT2 shows no detectable global conformational changes relative to the apo state: TM1b and TM6a remain unchanged, and the Arg86–Asp525 gate remains open, with Phe308 exhibiting the only structurally discernible local rotameric switch (**Fig. EV6E; Movie EV1**). This suggests that substrate binding initially induces local remodeling of the S1 pocket rather than global gate closure.

In contrast to the canonical LeuT-fold mechanism, we therefore propose that the observed structures represent an early substrate-bound intermediate preceding occluded states. At this stage, the rotameric switch of Phe308 reshapes the geometry and hydrophobic environment of the S1 pocket to enable substrate capture and may facilitate subsequent extracellular gating rearrangements. Collectively, Phe308 acts as a key early conformational sensor that links substrate recognition in the S1 pocket to downstream extracellular gating rearrangements during the transport cycle.

### Allosteric inhibition by LOR stabilizing the outward-occluded state of B^0^AT2

LOR, a tricyclic H₁ antihistamine widely used for allergy treatment, was identified as the first characterized inhibitor of B^0^AT2 (**Fig. 3A**) (Cuboni et al., 2014). We determined the cryo-EM structure of the B^0^AT2^LOR^ complex at a resolution of 2.74 Å (**Fig. EV2A–E**). In this structure, both the extracellular and intracellular vestibules are closed and a prominent T-shaped density corresponding to LOR is located above the S1 site on the extracellular side, occupying the previously identified allosteric S2 pocket (**Fig. 3B–C**) (Shi, Quick et al., 2008, Singh, Yamashita et al., 2007). This binding mode suggests that LOR stabilizes B^0^AT2 in an outward-occluded conformation. LOR, comprising a chlorinated tricyclic core, a piperidine ring, and an ethyl carbamate substituent (**Fig 3A**), is accommodated within the S2 pocket formed by TM1, TM3, TM6, TM8, TM10, and EL4 (**Fig. 3B–C**). At the binding site, the ethyl carbamate group of LOR extends toward the extracellular vestibule and forms an electrostatic contact with the side chain of Arg86 (**Fig. 3D–E**). By contrast, the piperidine and tricyclic rings are deeply inserted into a highly hydrophobic cavity lined by a battery of hydrophobic residues (**Fig. 3E–F**). The chlorine atom is positioned in proximity to Ser165, mediating polar interactions with the S2 pocket (**Fig. 3D–E**). To examine the importance of these interactions in LOR binding, we generated alanine substitutions of residues lining the S2 pocket (**Fig. 3G**). Our results showed that simultaneous alanine substitution of three hydrophobic residues (W85A/F463A/L467A) could effectively weaken LOR binding (**Fig. 3G; Fig. EV7A–C**). Notably, these mutations did not interfere with the expression, localization, and substrate transport activity of B^0^AT2 (**Fig. 3G; Fig. EV7A–D**), suggesting that LOR binds to the S2 pocket and inhibits B^0^AT2 activity through an allosteric mechanism.

**Figure 3.**
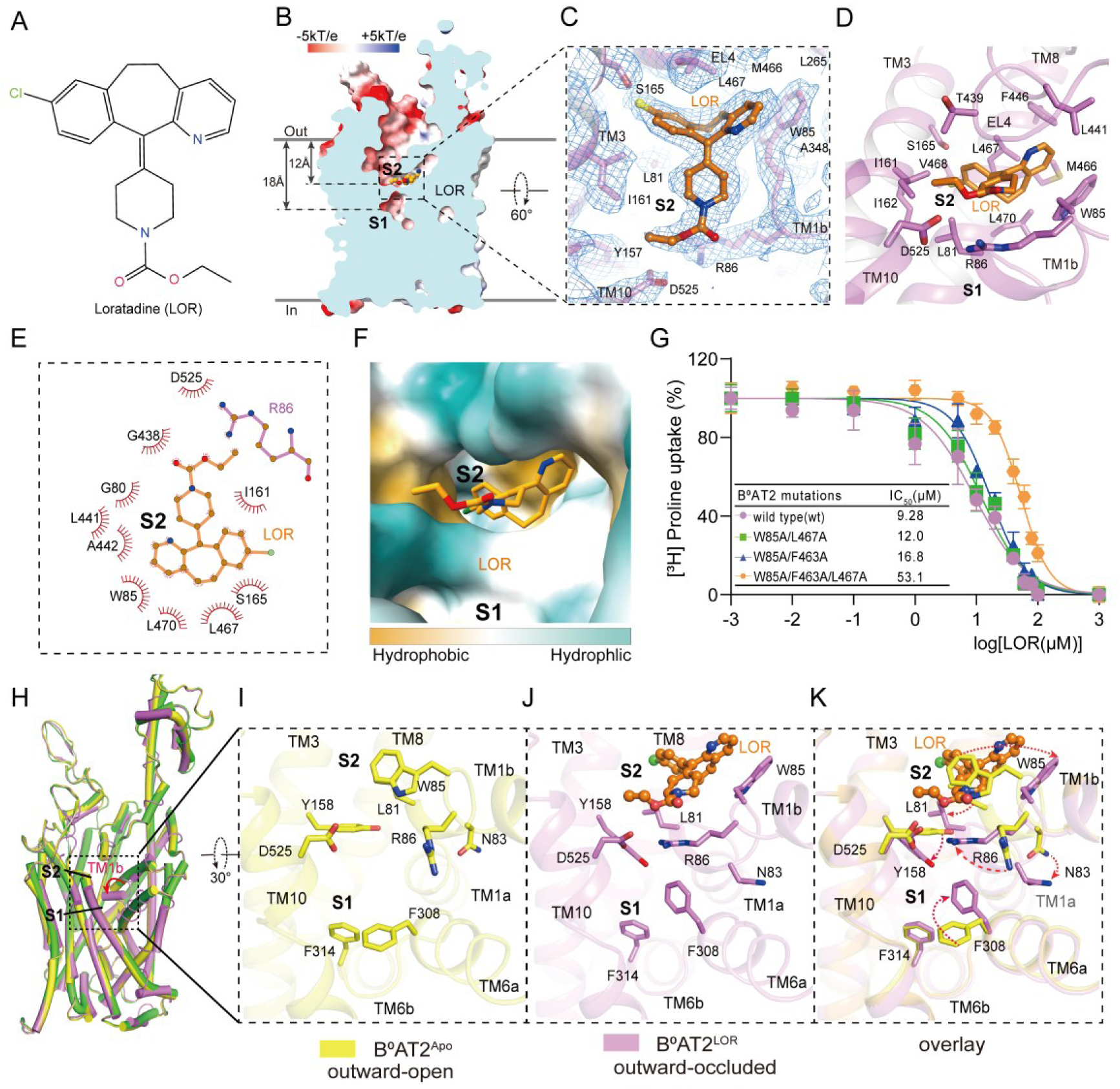
Structural basis of LOR-mediated allosteric inhibition of B^0^AT2 in the outward-occluded state. (A). Chemical structure of LOR. (B). Sagittal slice through an electrostatic surface potential of B^0^AT2^LOR^, with the S1 and S2 pockets labeled. Distances from the extracellular membrane are indicated. (**C**) Cryo-EM density (blue mesh) and model of LOR (orange sticks) in S2 pocket. (**D**). Interaction of LOR in S2 pocket and the key residues for interaction are shown as sticks. (E) Schematic representation of LOR binding interactions generated using LigPlot+, with hydrophobic interactions shown as red spokes. (F) Side view of LOR bound within the hydrophobic S2 pocket. Green and yellow indicate hydrophilic and hydrophobic regions, respectively. (G) Effects of mutations on LOR inhibitory potency measured by [³H]-proline uptake. Data are shown as means ± s.d.; n = 3 technical replicates. The experiment was performed independently twice with similar results. (**H-K**). Structural comparison of B^0^AT2^LOR^ (violet) and B^0^AT2^Apo^ (yellow). Regions undergoing conformational changes are boxed, with red arrows indicating displacement of the TM1b N-terminal segment (H). Close-up views of corresponding residues in the S1 (I) and S2 (J) pockets are shown, with LOR displayed as ball-and-stick models. (K) Superposition highlights residue rearrangements, with displacement directions indicated.

To further understand this inhibition mode, we superimposed B^0^AT2^LOR^ structure with that of B^0^AT2^Apo^ (**Fig. 3H–K; Movie EV2**). This comparison showed that LOR binding is associated with coordinated rearrangement in the extracellular gating elements, particularly TM1b and TM6a. Upon LOR binding, Trp85 rotates by ∼105°, enlarging the S2 cavity to accommodate the ligand (**Fig. 3K; Movie EV2**). Concurrently, Arg86 moves from the side of the extracellular vestibule toward its center and forms a salt bridge with Asp525, accompanied by local rearrangements of Phe308 and Tyr158 that cap the S1 pocket (**Fig. 3I–K**). Together, these changes reshape the S2 pocket and close the extracellular access channel to the S1 site. (**Fig. 3K; Movie EV2**). Thus, LOR stabilizes B^0^AT2 in an outward-occluded conformation by coupling S2 occupancy to closure of the S1–S2 gating network. This structural remodeling provides a mechanistic basis for LOR-mediated allosteric inhibition, in which substrate transport is blocked by preventing extracellular access to the S1 pocket.

Tricyclic and tricyclic-like antidepressants, including clomipramine (CMI) and maprotiline (Map), represent a class of LeuT-fold transporter ligands that share aromatic scaffolds with LOR and inhibit substrate uptake by blocking extracellular access (**Fig. EV8A**)(Singh et al., 2007, Zhang, Yin et al., 2024). To assess whether this binding mode is conserved across LeuT-fold transporters, we compared the B^0^AT2^LOR^ structure with representative structures of related ligand-bound LeuT-fold transporters, including NET bound to maprotiline (NET^Map^; PDB: 8Y8Z) and LeuT bound to clomipramine (LeuT^CMI^; PDB: 2Q6H) (**Fig. EV8B–D**)(Singh et al., 2007, Zhang et al., 2024). Despite similar chemical scaffolds, these ligands adopt distinct binding modes and stabilize different conformational states (**Fig. EV8B–D**). Specifically, Map binds the S1 site in NET to stabilize an outward-open state (Zhang et al., 2024), whereas CMI primarily engages the peripheral S2 region in LeuT and, together with an additional ligand molecule, stabilizes an outward-occluded state (Singh et al., 2007) (**Fig. EV8E–F**). In contrast, LOR is deeply embedded in the S2 pocket of B^0^AT2, where it reinforces gate closure and stabilizes an outward-occluded state through an allosteric mechanism (**Fig. EV8E–F**). Collectively, these structural comparisons reveal distinct ligand-binding modes across LeuT-fold transporters and highlight the structural variability of the extracellular vestibule, including the S2 site of B^0^AT2.

### Cooperative multi-site inhibition by TGI stabilizing the inward-open state of B^0^AT2

TGI is a nipecotic acid derivative containing a piperidine-3-carboxylate headgroup and two thienyl substituents (**Fig. 4A**). Although originally developed as a GAT1 inhibitor, it has been shown that TGI could also inhibit B^0^AT2. However, it is still not clear how TGI binds and inhibits B^0^AT2 (Kukulowicz et al., 2025). To address this question, we determined the cryo-EM structure of the B^0^AT2^TGI^ complex at a resolution of 3.51 Å (**Fig. EV2F–J**). The map shows that B^0^AT2 adopts an inward-open conformation (**Fig. 4B, D**). The map also reveals three non-protein densities on the intracellular side that could be well fitted by TGI molecules, which hereafter will be referred to as TGI-1, TGI-2, and TGI-3 (**Fig. 4B–C; Fig. EV9A–C; Movie EV3**). TGI-1 occupies the canonical S1 pocket (**Fig. 4D; Fig. EV9A**), whereas TGI-2 and TGI-3 reside in two adjacent intracellular cavities (**Fig 4C, E–F; Fig. EV9B–C**). TGI-2 is positioned between TM1a and TM7, and TGI-3 lies near the cytoplasmic end of TM6b adjacent to IL1 (**Fig. 4E–F; Fig. EV9B–C; Movie EV3**). We name these intracellular cavities as the S3 and S4 pockets, respectively (**Fig. 4B–C**).

**Figure 4.**
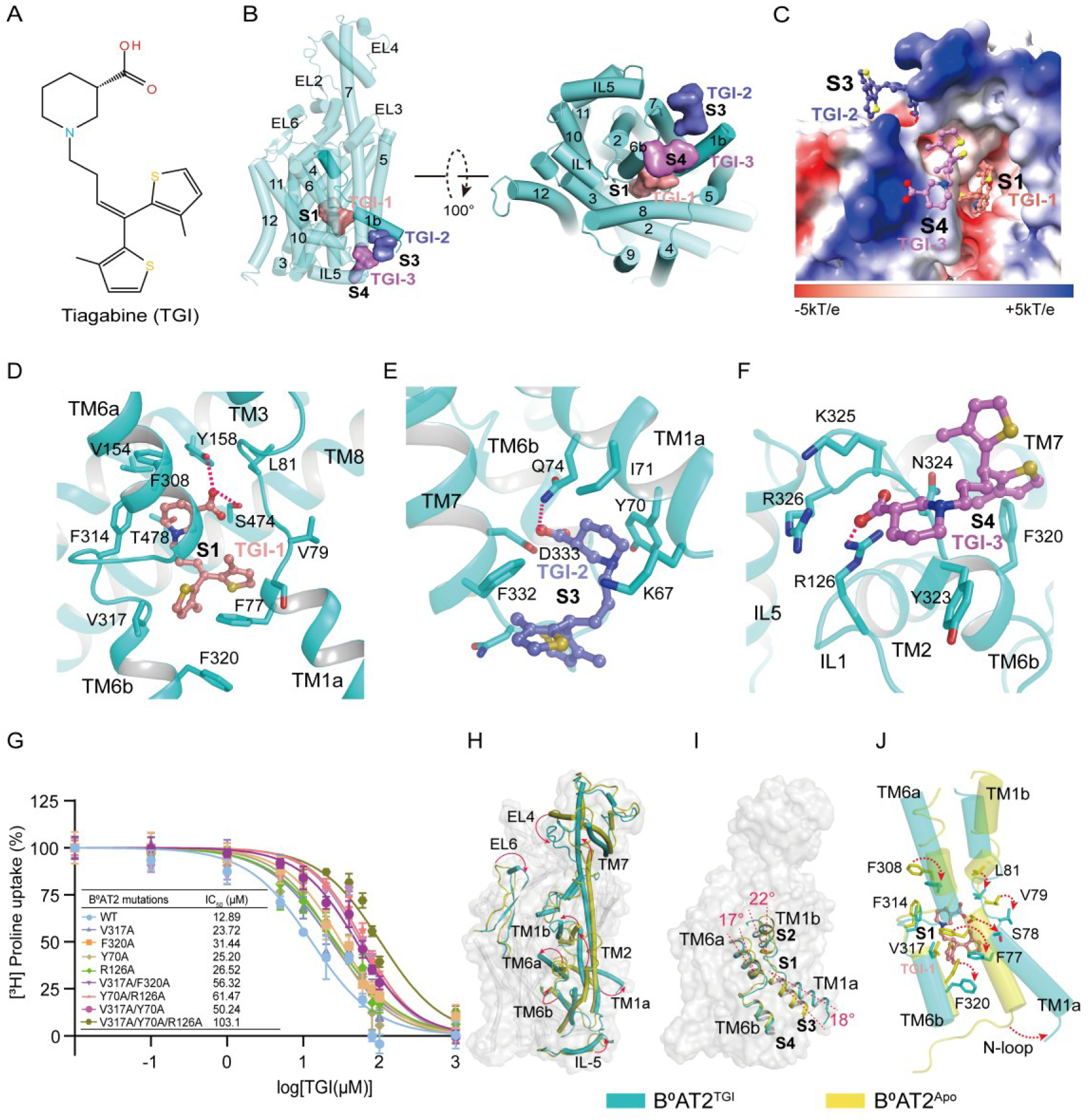
Structural basis of TGI-mediated cooperative multi-site inhibition of B^0^AT2 in the inward-open state. (A). Chemical structure of TGI. (B). Side and bottom views of the overall structure of B^0^AT2^TGI^ are shown in cylindrical representation. TGI-1, TGI-2, and TGI-3 are displayed as surface representations and colored salmon, slate, and violet, respectively. The transmembrane helices and the binding pockets S1, S3, and S4 are indicated. (C). Electrostatic potential surface of the B^0^AT2^TGI^ complex viewed from the cytoplasmic side. TGI-1, TGI-2, and TGI-3 are shown in ball-and-stick representation, occupying the S1, S3, and S4 pockets, respectively. (**D-F**) Interactions of TGI-1, TGI-2, and TGI-3 within the S1, S3, and S4 pockets, respectively, shown from different viewing orientations. Key interacting residues are depicted as sticks, and hydrogen bonds are indicated by dashed lines. (G) Effects of single mutations and S1/S3 (V317A/Y70A), S1/S4 (V317A/F320A), S3/S4 (Y70A/R126A), and S1/S3/S4 (V317A/Y70A/R126A) mutations on TGI inhibitory potency measured by [³H]-proline uptake. Data are shown as means ± s.d.; n = 3 technical replicates. The experiment was performed independently twice with similar results. (H) Structural comparison of the inward-open B^0^AT2^TGI^ (cyan) and outward-open B^0^AT2^Apo^ (yellow) conformations. Major conformational changes are indicated by red arrows. (I) Superposition of B^0^AT2^TGI^ and B^0^AT2^Apo^ highlighting tilting of TM1a, TM1b, and TM6a. TGI-1, TGI-2, and TGI-3 are shown as sticks. (J) Structural superposition of B^0^AT2^TGI^ in the inward-open state and B^0^AT2^Apo^ in the outward-open state. TGI-1 and key residues in the S1 binding pocket are shown as sticks. Residues involved in conformational changes are highlighted with red arrows.

TGI engages B^0^AT2 through an extensive network of polar, aromatic, and hydrophobic interactions. At S1, the TGI-1 carboxylate forms hydrogen bonds with Tyr158 and Ser474, whereas the piperidine and thienyl groups are stabilized by hydrophobic packing within a surrounding aromatic-aliphatic pocket (**Fig. 4D**). This binding mode resembles the orthosteric interactions observed for TGI in GAT1 and SLC6A20 (Broer et al., 2024, Motiwala, Aduri et al., 2022, Zhu, Huang et al., 2023), where the S1 pocket is expanded, leading to an upward displacement of TM1a (**Fig. EV9D–E**). TGI-2 in S3 is anchored by polar contacts involving Asp333 and Gln74 and packs against Leu67, Tyr70, and Phe332 (**Fig. 4E**), whereas TGI-3 in S4 engages Arg126, Arg326, and Asn324 and is stabilized by hydrophobic contacts involving Phe320 and Tyr323 (**Fig. 4F**). Together, these interactions position three TGI molecules into distinct pockets, where they likely act cooperatively to stabilize the inward-open state by reinforcing the displaced TM1a conformation.

To validate this cooperative multi-site inhibition mechanism, we generated single and combinatorial mutations of residues lining the S1, S3, and S4 pockets (**Fig. 4G**). Consistent with previous reports, TGI inhibits B^0^AT2 with an IC₅₀ of ∼12.89 μM (**Fig. 4G**) (Kukulowicz et al., 2025). Mutations targeting individual sites resulted in only modest decreases in inhibitor sensitivity (**Fig. 4G**), Notably, simultaneous substitutions across two or three pockets led to a progressively greater loss of TGI potency, indicating an additive effect on de-repression of inhibition (**Fig. 4G**). Importantly, these mutations did not affect B^0^AT2 expression, localization, and substrate transport activity compared with WT B^0^AT2 (**Fig. EV10A–D**). Collectively, these results establish a cooperative multi-site mechanism underlying TGI-mediated inhibition of B^0^AT2 involving the S1, S3, and S4 pockets.

To further understand the structural basis of this cooperative multi-site inhibition, we performed a comparative analysis of B^0^AT2^TGI^ with the outward-open B^0^AT2^Apo^ and outward-occluded B^0^AT2^LOR^ structures (**Fig. 4H–I; Fig. EV11A-E**). Notably, relative to B^0^AT2^Apo^, the B^0^AT2^TGI^ complex exhibits pronounced structural rearrangements involving TM1a, TM1b, TM6a, IL5, EL4, and EL6 (**Fig. 4H-I**), with TM1b and TM6a showing particularly convergent movements (**Fig. 4I; Movie EV3**). This is accompanied by tighter closure of the extracellular gate mediated by conserved gating residues, thereby preventing substrate translocation (**Fig. EV11A–C**). Concurrently, TM1a tilts upward by ∼18°, leading to expansion of the S1 pocket and exposure of the intracellular access channel (**Fig. 4I–J**). Importantly, the N-loop (residues 59–64), which connects to TM1a, is unresolved in the B^0^AT2^TGI^ structure, whereas in the B^0^AT2^Apo^ and B^0^AT2^LOR^ structures, the N-loop forms a stabilizing network centered on Arg61, Trp64, and Lys67 that contributes to the closure of the intracellular vestibule (**Fig. EV11D–E**). In line with previous reports, the N-loop has been reported to play an important role in conformational transitions and substrate transport in LeuT-fold transporters (Cheng & Bahar, 2014, Gotfryd, Boesen et al., 2020, Khan, Sohail et al., 2020, Wei et al., 2024). Interestingly, in B^0^AT2^TGI^ structure, TGI-3, acting as a steric blocker at S4, perturbs the Arg61-Asp488 and Trp64-Ser322 network via interactions with Arg126, whereas TGI-2, acting as a wedge at S3, disrupts Lys67-Asn328/Asp333/Tyr70 contacts and inserts between TM1a and TM7, collectively preventing intracellular gate resetting (**Fig. EV11C, E; Movie EV3**). Conservation and mutational analyses indicate that N-loop residues are essential for transport activity (**Fig. EV3 and EV11F**), supporting a role in which the N-loop functions as a key intracellular gating module that controls TM1a dynamics. Together, these findings reveal a mechanistic basis for cooperative multi-site inhibition, in which cooperative engagement of S1, S3, and S4 stabilizes the inward-open state by coupling TM1a displacement to disruption of the N-loop-mediated gate.

### S3 and S4 are conserved intracellular cavities with potential for modulation across the SLC6 family

To determine whether the newly identified intracellular pockets are unique to B^0^AT2 or represent conserved structural features of the SLC6 family, we compared B^0^AT2^TGI^ with multiple inward-open NSS transporter structures, including GAT1^TGI^ (PDB: 7Y7Z), SERT^HJM^ (PDB: 6DZZ), GlyT1^ALX^ (PDB: 8WFJ), and DAT1^GBR^ (PDB: 8Y2F) (Coleman, Yang et al., 2019, Li et al., 2024, Wei et al., 2024, Zhu et al., 2023) (**Fig. EV12A–E**), and further analyzed sequence conservation across the SLC6 family (**Fig. EV12A–E and EV13A–L**). Despite substantial differences in ligand chemistry (**Fig. EV12A**), these inhibitors stabilize a common inward-open state across different LeuT-fold transporters, thereby revealing a similar intracellular architectural framework formed by TM1a, TM6b, IL1, and TM7 that enables direct comparison with the B^0^AT2 intracellular vestibule (**Fig. EV12B–E**).This region can be partitioned into sub-pockets analogous to S3 and S4, which exhibit conservation and accessibility comparable to the well-characterized and druggable S1 pocket, suggesting that these intracellular cavities may also serve as potential ligand-binding sites (**Fig. EV12B–E, EV13A–I**). However, sequence and structural variations within these subpockets suggest that their modes of ligand engagement are likely to differ among transporters (**Fig. EV12B–E, EV13G–L**).Such variation, exemplified by residues corresponding to Tyr70 (S3) and Arg126 (S4) in B^0^AT2, that are substituted by distinct aromatic or polar/non-polar residues (**Fig. EV12B, D–E, EV13K–L**), indicates that local chemical diversity within a conserved intracellular architecture could provide a structural basis for transporter selectivity. Collectively, these analyses identify S3 and S4 as conserved intracellular cavities across the SLC6 family, suggesting their potential as intracellular sites for transporter modulation.

## Discussion

B^0^AT2 is a Na⁺-dependent LeuT-fold transporter that mediates neutral amino acid uptake and maintains neuronal amino acid homeostasis (Kukulowicz et al., 2023, Santarelli et al., 2016). Its dysfunction is linked to depression and stress-associated disorders (Liu, Quast et al., 2013, Santarelli et al., 2015). Here, we resolved multiple conformational states of B^0^AT2, including apo outward-open, early substrate-bound intermediates, LOR-stabilized outward-occluded, and TGI-bound inward-open states (**Fig. 5A–D**), revealing a structural trajectory underlying substrate capture, gating transitions, and state-specific inhibition.

**Figure 5.**
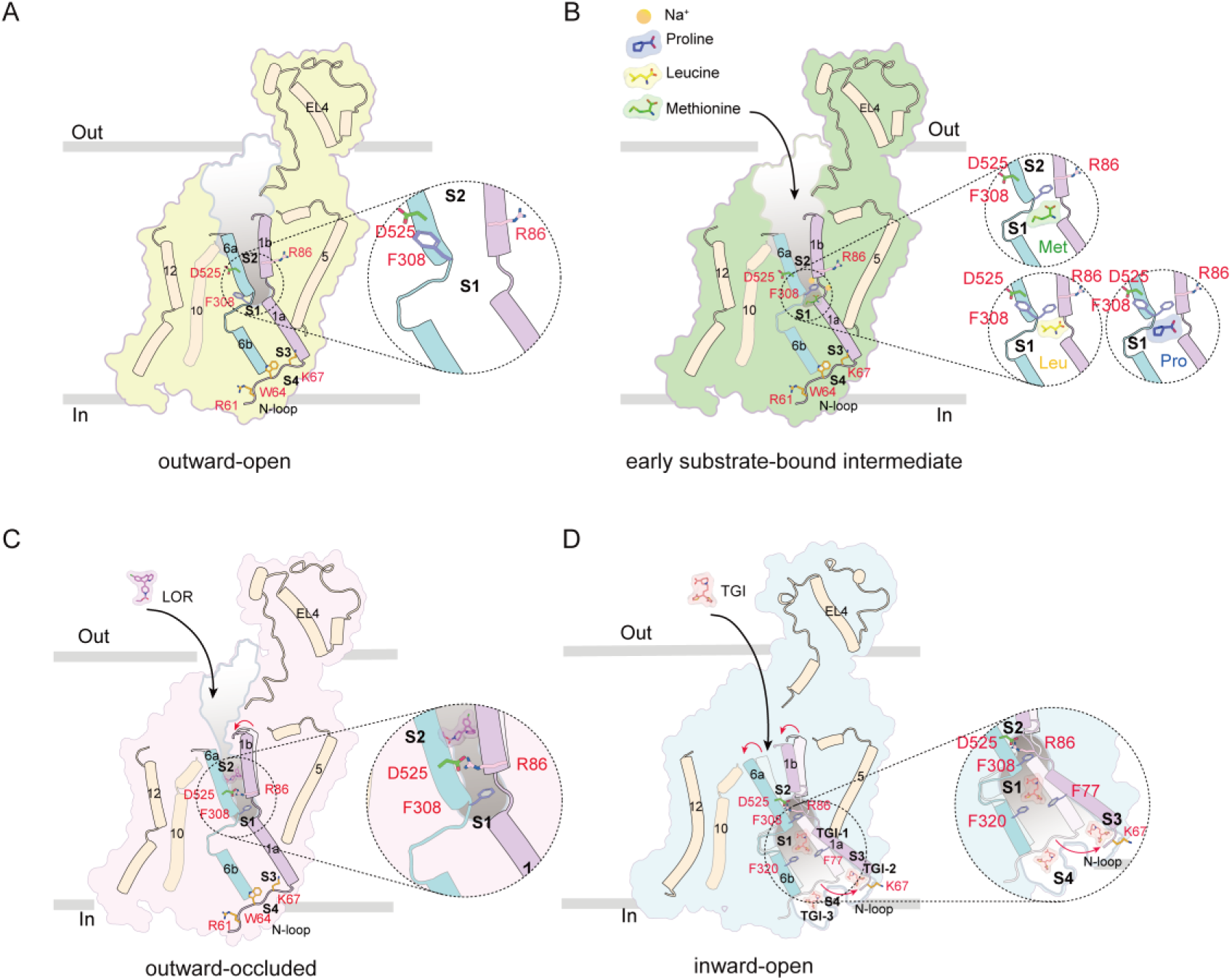
Substrate recognition and inhibitor mechanisms of B^0^AT2. (A) Structural overview of B^0^AT2 in the outward-open apo state. The dashed circle highlights the S1 pocket, which remains accessible to the extracellular vestibule. Key gating residues and transmembrane helices are indicated. (B) Structural overview of B^0^AT2 in early substrate-bound intermediate states. The dashed circle highlights the S1 pocket, illustrating that the side chain of Phe308 adopts distinct conformations depending on the bound substrate. Key gating residues and transmembrane helices are indicated. (C) Structural overview of B^0^AT2^LOR^ in an outward-occluded conformation, with LOR bound at the allosteric S2 site, where it seals the extracellular vestibule to block substrate access via allosteric inhibition. Key gating residues and transmembrane helices are indicated. (D) Structural overview of B^0^AT2 bound to three TGI molecules at the S1, S3, and S4 pockets. Their coordinated binding elevates TM1a and disrupts N-loop–mediated gating, stabilizing the inward-open state and enabling cooperative multi-site inhibition. Key gating residues and transmembrane helices are indicated.

Unlike canonical LeuT-fold transporters, where substrate binding is coupled to large-scale gating rearrangements, B^0^AT2 exhibits minimal global response upon substrate binding. Instead, a single residue, Phe308, undergoes a substrate-dependent rotamer switch that locally reshapes the S1 pocket. Notably, Phe308 also displays conformational heterogeneity across different substrate-bound states (Met, Leu, and Pro), suggesting a substrate-dependent energetic landscape underlying its rotamer selection (**Fig. 5A-B; Movie EV1**). This reveals a previously unappreciated mechanism in which early substrate recognition is governed by localized micro-switch tuning rather than global occlusion, positioning Phe308 as a primary determinant of substrate selectivity.

Tricyclic ligands acting at the extracellular vestibule further highlight the plasticity of the gating network. Although LOR, Map, and CMI share conserved aromatic scaffolds, they stabilize distinct conformational endpoints across LeuT-fold transporters. In B^0^AT2, LOR binds the S2 pocket and induces coupled rearrangements of TM1b, TM6a, and conserved gating residues, stabilizing an outward-occluded state (**Fig. 5C; Movie EV2**). Importantly, occlusion arises not from S1 blockade but from allosteric coupling between S2 occupancy and the S1–S2 gate, indicating an extracellular allosteric inhibition mechanism that reveals a highly adaptable vestibule capable of translating chemically similar ligands into divergent structural outcomes.

To date, most small-molecule inhibitors of LeuT-fold transporters target the conserved S1 or S2 pockets, whereas intracellular allosteric sites remain largely unexplored(Li et al., 2024, Singh et al., 2007, Wang, Goehring et al., 2013, Wei et al., 2024, Zhang et al., 2024). In this context, TGI reveals a fundamentally different inhibitory principle operating on the intracellular side. Rather than targeting a single orthosteric site, TGI engages a cooperative multi-site inhibition network of three spatially distinct cavities, stabilizing the inward-open state (**Fig. 5D; Movie EV3**). This multi-site mechanism couples S1 occupancy to two intracellular pockets exposed only in the inward-open conformation, locking the transporter through stabilization of TM1a displacement and disruption of intracellular gating architecture. These findings extend the mechanistic landscape of LeuT-fold inhibition beyond the classical S1/S2 paradigm and reveal that intracellular vestibules can function as active regulatory surfaces rather than passive solvent-exposed regions.

Importantly, comparative structural and sequence analyses across diverse LeuT-fold NSS transporters demonstrate that these intracellular cavities are not unique to B^0^AT2. Instead, they are embedded within a conserved architectural framework formed by TM1a, TM6b, IL1, and TM7. Within this conserved scaffold, subpockets corresponding to S3 and S4 exhibit both structural preservation and chemical variability across family members. This combination suggests a dual principle: a conserved geometric platform that maintains accessibility in the inward-open state, and a tunable chemical environment that may underlie transporter-specific ligand recognition. Such organization provides a structural basis for selective intracellular modulation and expands the druggable landscape of the SLC6 family beyond the well-characterized extracellular pockets.

Several limitations should be noted. First, cryo-EM structures represent discrete conformational states, and the kinetic coupling between these states remains to be elucidated. Second, although structural conservation suggests the presence of analogous intracellular cavities across SLC6 transporters, their physiological and pharmacological relevance remains to be experimentally validated.

Collectively, B^0^AT2 operates through distributed conformational micro-switches spanning both membrane sides: Phe308 tunes early substrate capture, S2-mediated allostery controls extracellular gating, and intracellular multi-site engagement locks the inward-open state. These findings redefine LeuT-fold transporter regulation as a multi-surface, micro-switch–driven process and identify intracellular vestibules as a structurally conserved yet chemically programmable platform for state-selective modulation.

## Methods

### Reagents and tools table

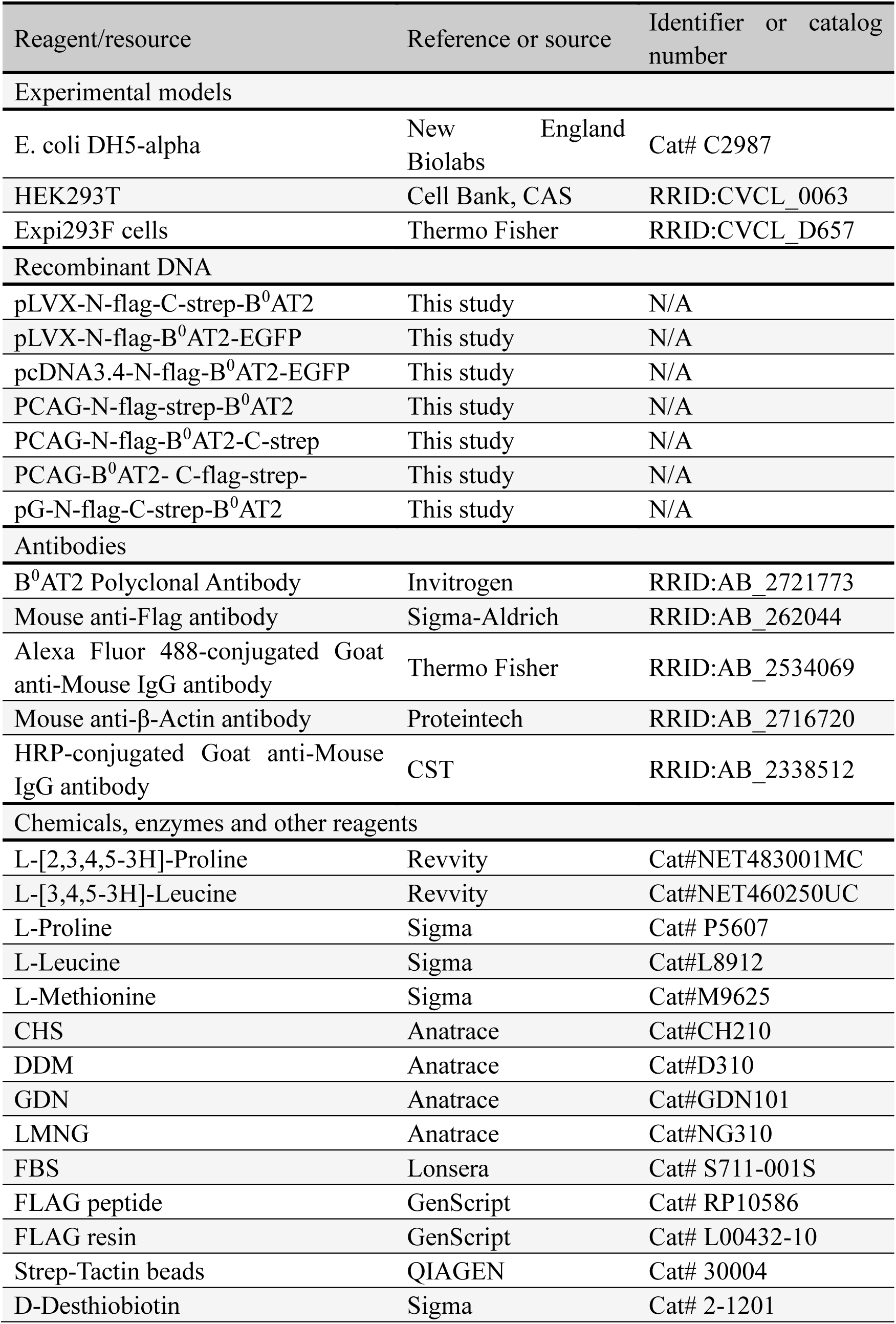
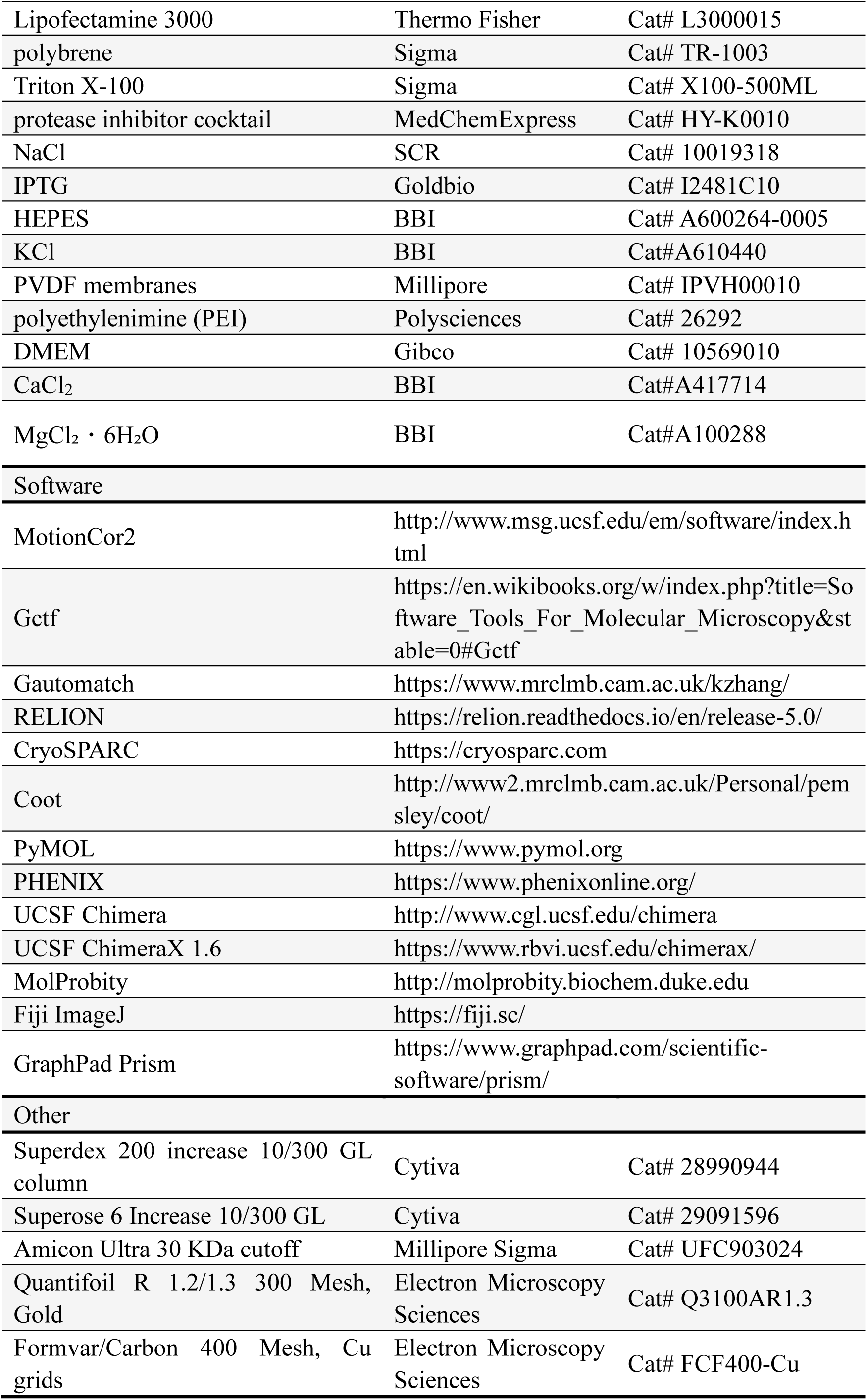

### Cell culture

The plasmids were amplified in E. coli Top10. HEK293T cells were obtained from ATCC (CRL-3216) and maintained in DMEM (Gibco, 10569010) supplemented with 10% FBS (Lonsera, S711-001S). Expi293F cells were obtained from Gibco (A14527) and cultured in Cell-Wise serum-free medium (CW001). All cells were maintained at 37 °C in a humidified incubator with 5% CO_2_.

### Plasmids and constructs

The human B^0^AT2 gene sequence (UniProt: Q9H2J7) was synthesized and cloned into a pCAG vector, containing an HRV-3C cleavage site, followed by Twin-Strep and Flag tags. Site-directed mutagenesis was performed by homologous recombination PCR, and all mutant constructs were verified by DNA sequencing (Shanghai Biosune Biotech Co., Ltd.). For expression screening, B^0^AT2 was also fused in frame to a C-terminal EGFP tag in pcDNA3.4, which enabled rapid assessment of expression and monodispersity by fluorescence-detection size-exclusion chromatography (FSEC) (Chen, McMullan et al., 2013).

### Expression and purification of human B^0^AT2

For transient expression of recombinant B^0^AT2, expi293F cells were grown at 37 °C with 5% CO_2_ to a density of approximately 2.5 × 10^6^ cells/mL. Plasmid DNA encoding B^0^AT2 (1 mg) and polyethyleneimine (PEI, 3 mg) were each diluted in 40 mL serum-free medium, incubated for 5 min at room temperature, combined, and incubated for an additional 15 min before addition to the culture. Twelve hours after transfection, sodium butyrate was added to a final concentration of 10 mM, and the cells were cultured for a further 60h. Cells were harvested by centrifugation at 1, 500g for 15 min, flash-frozen in liquid nitrogen, and stored at −80 °C for subsequent purification.

Protein purification was performed on ice or at 4 °C to maintain stability. Cells pellets were resuspended in Buffer A (25 mM HEPES, 150 mM NaCl, pH 7.4) supplemented with 1 mM phenylmethylsulfonyl fluoride (PMSF), 1 μg/mL aprotinin, 1 μg/mL leupeptin, and 1 μg/mL pepstatin. The cells were homogenized using a Dounce homogenizer with about 50 strokes. To solubilize the cell membrane, 1% (w/v) n-Dodecyl-β-D-maltoside (DDM, Anatrace) and 0.1% (w/v) cholesteryl hemisuccinate (CHS, Anatrace) were added to the suspension, followed by gentle mixing at 4 °C for 2 hours. Cell debris was removed by centrifugation at 35,000 rpm for 50 min at 4 °C. The resulting supernatant was further clarified by passing it through a 0.22 μm filter to eliminate any residual debris. The filtered supernatant was incubated with 4 mL pre-equilibrated StrepTactin Beads 4FF(QIAGEN) at 4 °C with gentle stirring for 2 hours. The beads were packed into a gravity column and initially washed with 10 column volumes (CV) of Buffer A supplemented with 1% lauryl maltose neopentyl glycol (LMNG, Anatrace) and 0.1% (w/v) CHS. In the following washes, the detergent concentration was gradually decreased to 0.003% (w/v) LMNG and 0.0003% (w/v) CHS, in three steps with 10 CV for each step. B^0^AT2 protein was eluted using Buffer A containing 5mM D-Desthiobiotin (Sigma), 0.005% (w/v) LMNG and 0.0005% (w/v) CHS. The eluted protein was further purified by incubation with anti-Flag G1 Affinity Resin (GenScript) overnight at 4 ℃, followed by elution using wash buffer containing 200 µg/mL 3× Flag peptide. Finally, the protein was subjected to size-exclusion chromatography (SEC) using a Superose 6 Increase 10/300 GL column (GE) equilibrated with Buffer B (25 mM HEPES, 150 mM NaCl, 0.003% (w/v) LMNG, 0.0003% (w/v) CHS, pH 7.4). Peak fractions from the SEC were pooled and concentrated to 10 or 12 mg/mL for cryo-EM grid preparation.

### Cryo-EM sample preparation and data collection

Negative staining was performed to evaluate the quality of the purified recombinant B^0^AT2 protein. Briefly, 4 µL of purified B^0^AT2 protein was applied onto glow-discharged copper grids supported by a thin carbon film for 25 seconds. The grids were then stained with a 2% (w/v) uranyl acetate solution at room temperature. Examination of the negatively stained grid was carried out using an FEI Talos L120C transmission electron microscope (Thermo Fisher Scientific) operated at 120 kV. For the preparation of B^0^AT2^Pro^, B^0^AT2^Leu^, and B^0^AT2^Met^ complexes, 200 µM substrates were maintained in all buffers throughout the purification process, and the purified proteins were subsequently incubated with 3 mM substrate prior to grid preparation. For the preparation of B^0^AT2^LOR^ and B^0^AT2^TGI^ complexes, purified B^0^AT2 protein was concentrated to ∼18 mg/mL and incubated with LOR or TGI at a 3–4-fold molar excess relative to the protein for 0.5 h before grid preparation. Protein samples were applied to glow-discharged Quantifoil Au R1.2/1.3 grids, blotted for 3s at 8 °C and 100% humidity, and vitrified in liquid ethane using an FEI Vitrobot Mark IV. Cryo-EM data were collected on a 300 kV FEI Titan Krios G3i microscope equipped with a K3 direct electron detector and an energy filter (20 eV slit width). For B^0^AT2^Apo^, B^0^AT2^Pro^, B^0^AT2^Leu^, B^0^AT2^Met^, B^0^AT2^LOR^, and B^0^AT2^TGI^, data were acquired at 81,000x magnification (pixel size: 1.1 Å) using EPU for automated acquisition. Each image stack was dose-fractionated into 32 frames over a 2.8 s exposure, with a total dose of ∼50 e⁻/Å² and a defocus range of −1.2 to −1.8 μm. The six datasets comprised 5,619, 6,373, 8,294, 9,884, 8,857, and 9,289 movies, respectively.

### Cryo-EM data processing

Movies were motion-corrected, dose-weighted, and binned using MotionCor2 (Zheng, Palovcak et al., 2017), and the contrast transfer function (CTF) of each micrograph was estimated with CTFFIND4 (Rohou & Grigorieff, 2015). Particles were picked using Gautomatch or Topaz (Bepler, Morin et al., 2019). For the B^0^AT2^Apo^ dataset, 10,329,738 particles were initially extracted with a pixel size of 2.2 Å and subjected to iterative 2D classification, Ab-Initio reconstruction, heterogeneous refinement, and 3D classification in cryoSPARC (Punjani, Rubinstein et al., 2017) and RELION (Scheres, 2012). The best class was re-extracted at 1.1 Å per pixel, followed by Non-uniform Refinement and focused 3D classification without alignment yielding a final 2.84 Å reconstruction from 186,351 particles (Table EV1). The B^0^AT2^Pro^, B^0^AT2^Leu^, B^0^AT2^Met^, B^0^AT2^LOR^, and B^0^AT2^TGI^ datasets were processed using a similar workflow and yielded final maps at 2.80 Å, 2.82 Å, 2.92 Å, 2.74 Å, and 3.51 Å, respectively (Table EV1).

### Model building and refinement

An AlphaFold (Jumper, Evans et al., 2021)-predicted model of B^0^AT2 (AF-Q9H2J7-F1) was used as the initial template. The model was rigid-body docked into the B^0^AT2^Apo^ cryo-EM map in ChimeraX (v.1.6) (Pettersen, Goddard et al., 2021), followed by iterative manual rebuilding in COOT (v.0.9.8) (Emsley, Lohkamp et al., 2010) and real-space refinement in PHENIX (v.1.19) (Adams, Afonine et al., 2010). The B^0^AT2^Pro^, B^0^AT2^Leu^, B^0^AT2^Met^, and B^0^AT2^LOR^ models were built using the same general procedure. The B^0^AT2^TGI^ model was built de novo against its corresponding cryo-EM density map. Geometry restraints for Pro, Leu, Met, LOR, and TGI were generated with eLBOW in PHENIX (Adams et al., 2010). Model quality was assessed using MolProbity (Chen, Arendall et al., 2010, Davis, Leaver-Fay et al., 2007), and side chains without well-defined density were trimmed before deposition. Final refinement statistics are summarized in Supplementary Table EV1. Structural figures were prepared using UCSF Chimera (Pettersen, Goddard et al., 2004), ChimeraX (Pettersen et al., 2021), and PyMOL (Pei & Grishin, 2014).

### Generation of the B^0^AT2-EGFP-HEK293T stable cell line

For the experiments in this study, a B^0^AT2-EGFP-HEK293T stable cell line was generated to enable inducible expression and facilitate visualization and biochemical analysis of B^0^AT2 and its mutants. HEK293T cells were cultured at 37 ℃ with 5% CO_2_ in DMEM supplemented with 10% FBS. Stable HEK293T cell lines expressing B^0^AT2 and its mutants fused with EGFP were generated using a lentivirus system following the manufacturer’s packaging protocol (pCW vector: pMD2G: pSPAX2 = 2:1:1). Lentivirus production was performed using Lipofectamine 3000 (Thermo Fisher Scientific) according to the manufacturer’s instructions. The cells were selected using puromycin at a final concentration of 5 μg/mL. For protein expression, stable cells were seeded on poly-D-lysine-coated six-well plates (Sangon Biotech) at a density of 1.0 × 10^6^ cells/mL, and doxycycline hydrochloride (1 μg/mL) was added 12 h after seeding to induce expression. Protein expression was verified by western blotting and fluorescence microscopy.

### [^3^H]- proline transport assay

For transport assays, either wild-type and mutant B^0^AT2-EGFP-HEK293T stable cell lines or transiently transfected cells were plated in poly-D-lysine-coated (Sangon Biotech) 96-well plates at a density of 2 × 10⁴ cells per well and cultured at 37 °C with 5% CO₂. Doxycycline hydrochloride (1 μg/mL) was added 30 hours prior to the assay to induce B^0^AT2 expression. On the day of the transport assay, the DMEM medium was removed, and the cells were washed twice with prewarmed Locke’s modified physiological solution (LCIS) buffer (20 mM HEPES, 140 mM NaCl, 2.5 mM KCl, 1.8 mM CaCl_2_, 1 mM MgCl_2_, pH 7.4) at 37 ℃ for equilibration. The uptake assay was carried out at room temperature by incubating the cells for 15 min in 100 μL of LCIS buffer containing 180 μM unlabeled proline (cold) and 0.5 μCi/mL ^3^H-proline (hot). To terminate the assay, the proline containing LCIS buffer was removed, and the cells were washed twice with 200 μL ice-cold LCIS buffer. The cells were lysed with 100 μL of 1% (w/v) sodium dodecyl sulfate (SDS), and the lysate was transferred to a scintillation counting plate containing 900 μL of scintillation cocktail (Liquiscint, Cat. LS-121). Radioactivity was measured using a Tri-carb 5110 TR Counter (Perkin Elmer, Waltham, USA).

For inhibitor experiments, cells were pre-incubated with LOR or TGI for 30 min, and then incubated for an additional 15 min in the presence of both inhibitor and radiolabeled substrate. Background signal was defined using 100 μM LOR or TGI. Specific uptake was calculated by subtracting background from total uptake, and normalizing to the signal measured in the absence of inhibitor. Data were obtained from at least three independent experiments. IC_50_ values were determined in GraphPad Prism 10 by nonlinear regression of inhibitor concentration-response curves.

### Confocal Image Analysis

To examine the subcellular localization of B^0^AT2, HEK293T cells were seeded at ∼50% confluency in poly-L-lysine–coated 20 mm glass-bottom dishes (NEST, 801001). The following day, WT B^0^AT2 construct with a C-terminal Flag tag (pCAG vector) and corresponding mutant constructs were transfected into HEK293T cells using Lipofectamine 3000 (Thermo Fisher Scientific, L3000015) according to the manufacturer’s instructions. Twenty-four hours post-transfection, cells were washed with PBS and fixed with 4% paraformaldehyde (Beyotime Biotechnology, P0099) for 15 min at room temperature. Cells were permeabilized with 0.1% Triton X-100 in PBS for 10 min and blocked with 5% BSA for 1 h at room temperature. Cells were incubated with mouse anti-Flag primary antibody (Sigma-Aldrich, F1804; 1:500) at 4 °C overnight, followed by incubation with Alexa Fluor 488–conjugated goat anti-mouse IgG secondary antibody (Thermo Fisher Scientific, A-11001; 1:1000) for 1 h at room temperature in the dark.

Before imaging, cells were treated with SlowFade Diamond Antifade reagent containing with 4′,6-diamidino-2-phenylindole (DAPI) (Thermo Fisher Scientific, S36968) and sealed with a coverslip. Images were acquired using a Leica SP8 confocal microscope equipped with a 63× oil immersion objective. Alexa Fluor 488 fluorescence was excited using a 488 nm laser line, and emission signals were collected using a hybrid detector (HyD). DAPI was excited using a 405 nm laser line and detected in the corresponding emission channel.

### Western blot

To detect expression, the WT B^0^AT2 construct with a C-terminal Flag tag (pCAG vector) and related mutant constructs were transfected into HEK293T cells using Lipofectamine 3000 according to the manufacturer’s instructions. Twenty-four hours post-transfection, cells were washed three times with ice-cold PBS and lysed in RIPA lysis buffer (50 mM Tris-HCl pH 7.4, 150 mM NaCl, 1% NP-40, 0.1% SDS, 0.5% sodium deoxycholate), supplemented with protease inhibitor cocktail (Roche, 04693132001) and phosphatase inhibitor cocktail (Roche, 4906845001). Lysates were incubated on ice for 30 min and centrifuged at 12,000 rpm for 10 min at 4 °C to remove insoluble debris. Protein concentrations were determined using a bicinchoninic acid (BCA) assay. Equal amounts of protein (∼30 μg) were mixed with SDS loading buffer and denatured by heating at 70 °C for 10 min. Samples were separated by SDS-PAGE and transferred onto polyvinylidene difluoride (PVDF) membranes (Merck Millipore, IPVH00010) at 300 mA for 90 min. Membranes were blocked with 5% non-fat milk in TBST buffer (20 mM Tris, 150 mM NaCl, 0.1% Tween-20, pH 7.6) and incubated overnight at 4 °C with mouse monoclonal primary antibodies against Flag (Sigma-Aldrich, F1804; 1:1000) or β-Actin (Proteintech, 66009-1-Ig; 1:5000). After washing three times with TBST, membranes were incubated with horseradish peroxidase (HRP)-conjugated goat anti-mouse IgG secondary antibody (CST, 7076S; 1:3000) for 1 h at room temperature. Protein bands were visualized using Immobilon Western HRP substrate (Millipore, WBKLS0050), and band intensities were quantified using ImageJ software (Schneider, Rasband et al., 2012).

### Statistical Analysis

Global resolution estimations of cryo-EM density maps are based on the Fourier shell correlation (FSC) 0.143 criterion (Chen et al., 2013) The local resolution was estimated using cryoSPARC (Punjani et al., 2017). The number of independent experiments (n), biological replicates, and relevant statistical parameters are indicated in the figure legends where applicable. Western blot quantification was performed using GraphPad Prism v10 (GraphPad Software). Data are presented as mean ± SEM. Statistical significance was assessed using unpaired two-tailed Student’s t-test or one-way ANOVA with appropriate post hoc tests, as indicated. A p value < 0.05 was considered statistically significant. Statistics of the cryo-EM data collection, refinement, and validation were calculated using RELION (Scheres, 2012), cryoSPARC (Punjani et al., 2017), PHENIX (Adams et al., 2010), MolProbity (Chen et al., 2010) and PyMOL (Pei & Grishin, 2014) software packages.

## Data availability

Atomic models have been deposited in the Protein Data Bank (PDB) and the corresponding cryo-EM maps have been deposited in the Electron Microscopy Data Bank (EMDB), with accession codes: B^0^AT2^Apo^ (PDB 22UJ, EMD-68684); B^0^AT2^Pro^ (PDB 22XA, EMD-68745); B^0^AT2^Leu^ (PDB 22WU, EMD-68740); B^0^AT2^Met^ (PDB 22VB, EMD-68705); B^0^AT2^LOR^ (PDB 22WA, EMD-68730); and B^0^AT2^TGI^ (PDB 22XL, EMD-68754). All data are publicly available as of the date of publication.

## Acknowledgements

We thank Yakun Liang, Haishuang Chang and staff at the cryo-EM center of Shanghai Institute of Precision Medicine for their assistance with microscope calibration and data collection. We thank Hong Lu, Shufang He, Jie Huang, Ying Cui, Rijing Liao, and Yan Cai from Shanghai Jiao Tong University School of Medicine (SHSMU) for help with confocal microscopy and mass spectrometry analyses. We thank Huahua Song and Jinghua Zhang for providing experimental instruments of Experimental Nuclear Medicine Laboratory, Core Facility of Basic Medical Sciences, Shanghai Jiao Tong University School of Medicine. Financial support was supported by grants from the National Natural Science Foundation of China (32471344 to P.L., 32471339 to J.W., 31930063 and 32430033 to M.Lei.), the Shanghai Municipal Education Commission Gaofeng Clinical Medicine Grant Support (20181711 to J.W.), the Innovative Research Team of High-level Local University in Shanghai (SHSMU-ZLCX20211700 to J.W. and M.Lei.), the National Research Center for Translational Medicine at Shanghai, Ruijin Hospital, Shanghai Jiao Tong University School of Medicine, Shanghai, China (NRCTM (SH) -2021-01 to M.Lei.). Ming Lei is a SANS Exploration Scholar.

## Author Contributions

Yunlei.C. and Yunlong.C. carried out protein biochemistry experiments and cryo-EM data acquisition with the help of Y.S., H.L. Q.Y. and M.C.; Yunlei.C. and Yunlong.C. performed the cryo-EM data analysis and reconstruction with the help S.L., F.W., M.Li., and S.H.; Q.W. and Yunlei.C. performed the biochemical and imaging experiments with the help S.S and C.Z.; J.W., D.Y., Yunlei.C. and Yunlong.C. built the atomic structural model; J.W., J.X., S.C. and W.X. provided extensive discussions; M.Lei., Yunlei.C., Yunlong.C., and P.L. wrote the manuscript with the input from all authors. M.Lei., Yunlei.C., and P.L initiated and orchestrated the project.

## References

Adams PD, Afonine PV, Bunkoczi G, Chen VB, Davis IW, Echols N, Headd JJ, Hung LW, Kapral GJ, Grosse-Kunstleve RW, McCoy AJ, Moriarty NW, Oeffner R, Read RJ, Richardson DC, Richardson JS, Terwilliger TC, Zwart PH (2010) PHENIX: a comprehensive Python-based system for macromolecular structure solution. Acta Crystallogr D Biol Crystallogr 66: 213–21

Alquier T, Drgonova J, Jacobsson JA, Han JC, Yanovski JA, Fredriksson R, Marcus C, Schiöth HB, Uhl GR (2013) Involvement of the Neutral Amino Acid Transporter SLC6A15 and Leucine in Obesity-Related Phenotypes. PLoS ONE 8

Bepler T, Morin A, Rapp M, Brasch J, Shapiro L, Noble AJ, Berger B (2019) Positive-unlabeled convolutional neural networks for particle picking in cryo-electron micrographs. Nature methods 16: 1153–1160

Broer A, Hu Z, Kukulowicz J, Yadav A, Zhang T, Dai L, Bajda M, Yan R, Broer S (2024) Cryo-EM structure of ACE2-SIT1 in complex with tiagabine. J Biol Chem 300: 107687

Bröer A, Tietze N, Kowalczuk S, Chubb S, Munzinger M, Bak Lasse K, Bröer S (2005) The orphan transporter v7-3 (slc6a15) is a Na+-dependent neutral amino acid transporter (B0AT2). Biochemical Journal 393: 421–430

Chandra R, Francis TC, Nam H, Riggs LM, Engeln M, Rudzinskas S, Konkalmatt P, Russo SJ, Turecki G, Iniguez SD, Lobo MK (2017) Reduced Slc6a15 in Nucleus Accumbens D2-Neurons Underlies Stress Susceptibility. The Journal of Neuroscience 37: 6527–6538

Chen S, McMullan G, Faruqi AR, Murshudov GN, Short JM, Scheres SH, Henderson R (2013) High-resolution noise substitution to measure overfitting and validate resolution in 3D structure determination by single particle electron cryomicroscopy. Ultramicroscopy 135: 24–35

Chen VB, Arendall WB, 3rd, Headd JJ, Keedy DA, Immormino RM, Kapral GJ, Murray LW, Richardson JS, Richardson DC (2010) MolProbity: all-atom structure validation for macromolecular crystallography. Acta Crystallogr D Biol Crystallogr 66: 12–21

Cheng MH, Bahar I (2014) Complete mapping of substrate translocation highlights the role of LeuT N-terminal segment in regulating transport cycle. PLoS computational biology 10: e1003879

Coleman JA, Green EM, Gouaux E (2016) X-ray structures and mechanism of the human serotonin transporter. Nature 532: 334–9

Coleman JA, Yang D, Zhao Z, Wen P-C, Yoshioka C, Tajkhorshid E, Gouaux E (2019) Serotonin transporter–ibogaine complexes illuminate mechanisms of inhibition and transport. Nature 569: 141–145

Cuboni S, Devigny C, Hoogeland B, Strasser A, Pomplun S, Hauger B, Höfner G, Wanner KT, Eder M, Buschauer A, Holsboer F, Hausch F (2014) Loratadine and Analogues: Discovery and Preliminary Structure–Activity Relationship of Inhibitors of the Amino Acid Transporter B0AT2. Journal of Medicinal Chemistry 57: 9473–9479

Cui L, Li S, Wang S, Wu X, Liu Y, Yu W, Wang Y, Tang Y, Xia M, Li B (2024) Major depressive disorder: hypothesis, mechanism, prevention and treatment. Signal Transduction and Targeted Therapy 9

Davis IW, Leaver-Fay A, Chen VB, Block JN, Kapral GJ, Wang X, Murray LW, Arendall WB, 3rd, Snoeyink J, Richardson JS, Richardson DC (2007) MolProbity: all-atom contacts and structure validation for proteins and nucleic acids. Nucleic Acids Res 35: W375–83

Emsley P, Lohkamp B, Scott WG, Cowtan K (2010) Features and development of Coot. Acta Crystallogr D Biol Crystallogr 66: 486–501

G.R. Uhl SK, P. Gregor, E. Nanthakumar, A. Persico and S. Shimada (1992) Neurotransmitter transporter family cDNAs in a rat midbrain library: ‘orphan transporters’ suggest sizable structural variations. Molecular Brain Research 16: 353–359

Gotfryd K, Boesen T, Mortensen JS, Khelashvili G, Quick M, Terry DS, Missel JW, LeVine MV, Gourdon P, Blanchard SC, Javitch JA, Weinstein H, Loland CJ, Nissen P, Gether U (2020) X-ray structure of LeuT in an inward-facing occluded conformation reveals mechanism of substrate release. Nat Commun 11: 1005

Jumper J, Evans R, Pritzel A, Green T, Figurnov M, Ronneberger O, Tunyasuvunakool K, Bates R, Zidek A, Potapenko A, Bridgland A, Meyer C, Kohl SAA, Ballard AJ, Cowie A, Romera-Paredes B, Nikolov S, Jain R, Adler J, Back T et al. (2021) Highly accurate protein structure prediction with AlphaFold. Nature 596: 583–589

Khan JA, Sohail A, Jayaraman K, Szöllősi D, Sandtner W, Sitte HH, Stockner T (2020) The Amino Terminus of LeuT Changes Conformation in an Environment Sensitive Manner. Neurochemical research 45: 1387–1398

Kohli Martin A, Lucae S, Saemann Philipp G, Schmidt Mathias V, Demirkan A, Hek K, Czamara D, Alexander M, Salyakina D, Ripke S, Hoehn D, Specht M, Menke A, Hennings J, Heck A, Wolf C, Ising M, Schreiber S, Czisch M, Müller Marianne B et al. (2011) The Neuronal Transporter Gene SLC6A15 Confers Risk to Major Depression. Neuron 70: 252–265

Kristensen AS, Andersen J, Jorgensen TN, Sorensen L, Eriksen J, Loland CJ, Stromgaard K, Gether U (2011) SLC6 neurotransmitter transporters: structure, function, and regulation. Pharmacol Rev 63: 585–640

Kukulowicz J, Pietrzak-Lichwa K, Klimonczyk K, Idlin N, Bajda M (2023) The SLC6A15-SLC6A20 Neutral Amino Acid Transporter Subfamily: Functions, Diseases, and Their Therapeutic Relevance. Pharmacol Rev 76: 142–193

Kukulowicz J, Siwek A, Wolak M, Broer A, Yadav A, Broer S, Bajda M (2025) Insight into the Structure of the Neutral Amino Acid Transporter B(0)AT2 Enabled the Discovery of Tiagabine as an Inhibitor. ACS Chem Neurosci 16: 262–274

Li Y, Wang X, Meng Y, Hu T, Zhao J, Li R, Bai Q, Yuan P, Han J, Hao K, Wei Y, Qiu Y, Li N, Zhao Y (2024) Dopamine reuptake and inhibitory mechanisms in human dopamine transporter. Nature

Liu C, Quast C, Cuboni S, Bader D, Altmann A, Weber P, Arloth J, Röh S, Brückl T, Ising M, Kopczak A, Erhardt A, Hausch F, Lucae S, Binder EB (2013) Functional Coding Variants in SLC6A15, a Possible Risk Gene for Major Depression. PLoS ONE 8

Motiwala Z, Aduri NG, Shaye H, Han GW, Lam JH, Katritch V, Cherezov V, Gati C (2022) Structural basis of GABA reuptake inhibition. Nature 606: 820–826

Navratna V, Gouaux E (2019) Insights into the mechanism and pharmacology of neurotransmitter sodium symporters. Curr Opin Struct Biol 54: 161–170

Nyola A, Karpowich NK, Zhen J, Marden J, Reith ME, Wang DN (2010) Substrate and drug binding sites in LeuT. Curr Opin Struct Biol 20: 415–22

Pei J, Grishin NV (2014) PROMALS3D: multiple protein sequence alignment enhanced with evolutionary and three-dimensional structural information. Methods Mol Biol 1079: 263–71

Penmatsa A, Gouaux E (2014) How LeuT shapes our understanding of the mechanisms of sodium-coupled neurotransmitter transporters. J Physiol 592: 863–9

Pettersen EF, Goddard TD, Huang CC, Couch GS, Greenblatt DM, Meng EC, Ferrin TE (2004) UCSF Chimera--a visualization system for exploratory research and analysis. J Comput Chem 25: 1605–12

Pettersen EF, Goddard TD, Huang CC, Meng EC, Couch GS, Croll TI, Morris JH, Ferrin TE (2021) UCSF ChimeraX: Structure visualization for researchers, educators, and developers. Protein Sci 30: 70–82

Pramod AB, Foster J, Carvelli L, Henry LK (2013) SLC6 transporters: structure, function, regulation, disease association and therapeutics. Mol Aspects Med 34: 197–219

Punjani A, Rubinstein JL, Fleet DJ, Brubaker MA (2017) cryoSPARC: algorithms for rapid unsupervised cryo-EM structure determination. Nature methods 14: 290–296

Rohou A, Grigorieff N (2015) CTFFIND4: Fast and accurate defocus estimation from electron micrographs. J Struct Biol 192: 216–21

Santarelli S, Namendorf C, Anderzhanova E, Gerlach T, Bedenk B, Kaltwasser S, Wagner K, Labermaier C, Reichel J, Drgonova J, Czisch M, Uhr M, Schmidt MV (2015) The amino acid transporter SLC6A15 is a regulator of hippocampal neurochemistry and behavior. Journal of Psychiatric Research 68: 261–269

Santarelli S, Wagner KV, Labermaier C, Uribe A, Dournes C, Balsevich G, Hartmann J, Masana M, Holsboer F, Chen A, Muller MB, Schmidt MV (2016) SLC6A15, a novel stress vulnerability candidate, modulates anxiety and depressive-like behavior: involvement of the glutamatergic system. Stress 19: 83–90

Scheres SH (2012) RELION: implementation of a Bayesian approach to cryo-EM structure determination. J Struct Biol 180: 519–30

Schneider CA, Rasband WS, Eliceiri KW (2012) NIH Image to ImageJ: 25 years of image analysis. Nature methods 9: 671–5

Shi L, Quick M, Zhao Y, Weinstein H, Javitch JA (2008) The mechanism of a neurotransmitter:sodium symporter--inward release of Na+ and substrate is triggered by substrate in a second binding site. Mol Cell 30: 667–77

Singh SK, Yamashita A, Gouaux E (2007) Antidepressant binding site in a bacterial homologue of neurotransmitter transporters. Nature 448: 952–6

Szopa A, Poleszak E, Doboszewska U, Herbet M, Świąder K, Wyska E, Serefko A, Wlaź A, Korga A, Ostrowska M, Juś P, Jedynak S, Dudka J, Wlaź P (2018) Withdrawal of caffeine after its chronic administration modifies the antidepressant-like activity of atypical antidepressants in mice. Changes in cortical expression of Comt, Slc6a15 and Adora1 genes. Psychopharmacology 235: 2423–2434

Takanaga H, Mackenzie B, Peng J-B, Hediger MA (2005) Characterization of a branched-chain amino-acid transporter SBAT1 (SLC6A15) that is expressed in human brain. Biochemical and Biophysical Research Communications 337: 892–900

Ulrich H, Hägglund MGA, Roshanbin S, Löfqvist E, Hellsten SV, Nilsson VCO, Todkar A, Zhu Y, Stephansson O, Drgonova J, Uhl GR, Schiöth HB, Fredriksson R (2013) B0AT2 (SLC6A15) Is Localized to Neurons and Astrocytes, and Is Involved in Mediating the Effect of Leucine in the Brain. PLoS ONE 8

Wang H, Goehring A, Wang KH, Penmatsa A, Ressler R, Gouaux E (2013) Structural basis for action by diverse antidepressants on biogenic amine transporters. Nature 503: 141–5

Wang KH, Penmatsa A, Gouaux E (2015) Neurotransmitter and psychostimulant recognition by the dopamine transporter. Nature 521: 322–7

Wang L, Liu Z, Cao X, Li J, Zhang A, Sun N, Yang C, Zhang K (2017) A Combined Study of SLC6A15 Gene Polymorphism and the Resting-State Functional Magnetic Resonance Imaging in First-Episode Drug-Naive Major Depressive Disorder. Genetic Testing and Molecular Biomarkers 21: 523–530

Wei Y, Li R, Meng Y, Hu T, Zhao J, Gao Y, Bai Q, Li N, Zhao Y (2024) Transport mechanism and pharmacology of the human GlyT1. Cell 187: 1719–1732 e14

Yamashita A, Singh SK, Kawate T, Jin Y, Gouaux E (2005) Crystal structure of a bacterial homologue of Na+/Cl--dependent neurotransmitter transporters. Nature 437: 215–23

Zhang H, Yin Y-L, Dai A, Zhang T, Zhang C, Wu C, Hu W, He X, Pan B, Jin S, Yuan Q, Wang M-W, Yang D, Xu HE, Jiang Y (2024) Dimerization and antidepressant recognition at noradrenaline transporter. Nature

Zheng SQ, Palovcak E, Armache JP, Verba KA, Cheng Y, Agard DA (2017) MotionCor2: anisotropic correction of beam-induced motion for improved cryo-electron microscopy. Nature methods 14: 331–332

Zhu A, Huang J, Kong F, Tan J, Lei J, Yuan Y, Yan C (2023) Molecular basis for substrate recognition and transport of human GABA transporter GAT1. Nature Structural & Molecular Biology 30: 1012–1022

