## Supplementary Information for "Structural basis of substrate recognition and allosteric inhibition in human B^0^AT2"

**Table of Contents** **Page No.**

### Expanded View Figures

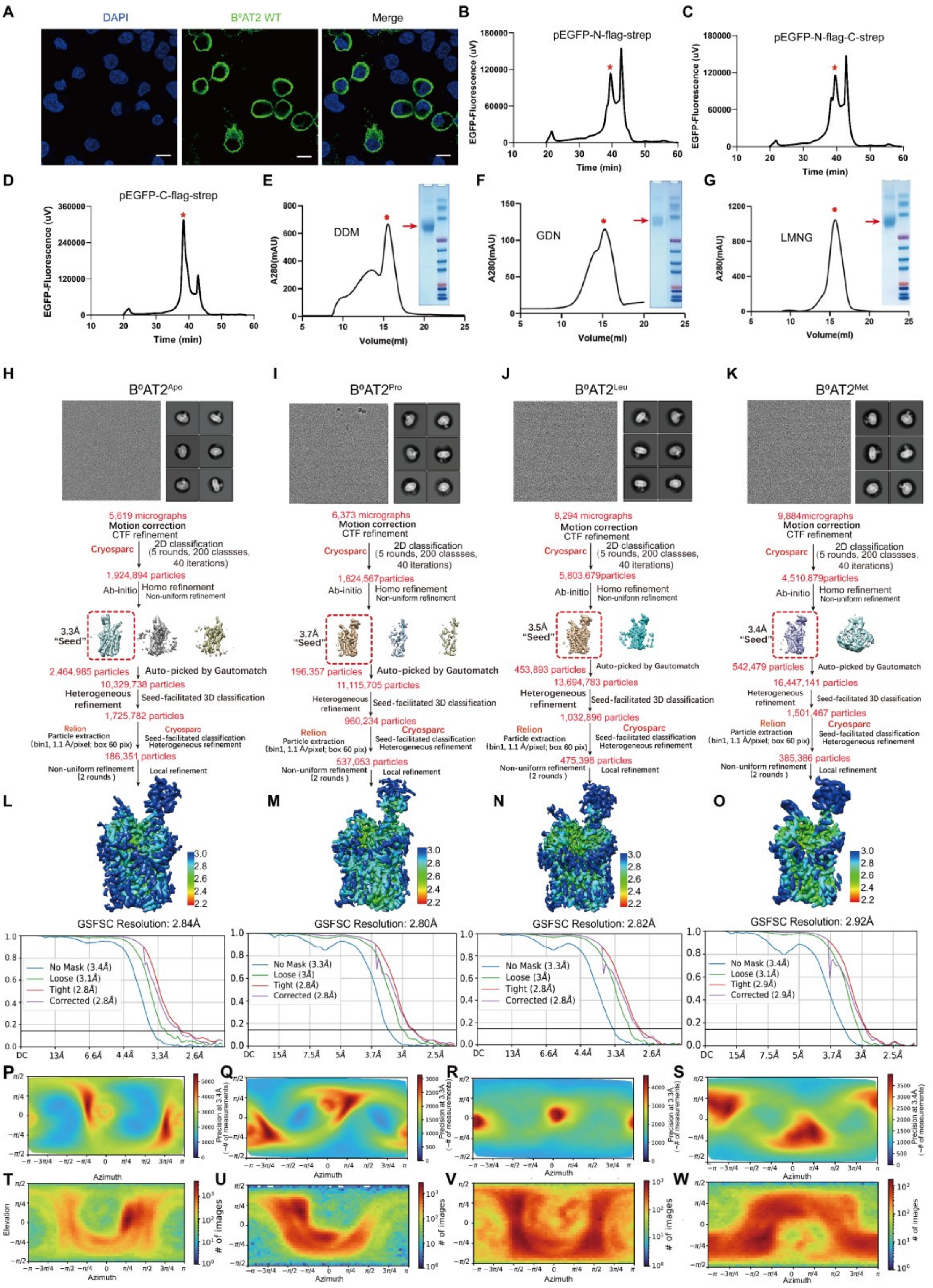

**Figure EV1. Functional characterization, purification, and cryo-EM data processing of B$^0$AT2.**

(**A**) Representative images of anti-FLAG immunostaining in B$^0$AT2-WT and mutants transfected HEK-293T cells. Scale bar = 5 μm.

(**B-D**) The well-behaved $B^0AT2$ constructs assayed by FSEC technology (Kawate & Gouaux, 2006). Fluorescence chromatogram of the construct pEGFP-N-flag-strep (B), pEGFP-N-flag-C-strep (C) and pEGFP-C-flag-strep (D) detected by FSEC technology, respectively. The red hexagon represents the peak of the fusion protein.

(**E–G**) Representative profiles of $B^0AT2$ in DDM (E), GDN (F), or LMNG (G) by size-exclusion chromatography (Superose 6 Increase) and SDS–PAGE analysis of $B^0AT2$ samples stained with Coomassie Blue. The red hexagon indicates the peak of the fusion protein. Peak fractions of $B^0AT2$ in LMNG were pooled and concentrated for cryo-EM studies. Experiments were repeated at least three times with similar results.

(**H–W**) Flowchart of cryo-EM data processing for $B^0AT2^{Apo}$, $B^0AT2^{Pro}$, $B^0AT2^{Leu}$ and $B^0AT2^{Met}$, showing final maps colored by local resolution and particle angular distribution. Representative raw micrographs and 2D class averages reveal distinct secondary structure features. Particle cleaning was performed through several rounds of classification, followed by CTF refinement and local refinement. Final maps were obtained at 2.84 Å, 2.80 Å, 2.82 Å and 2.92 Å according to the GSFSC (gold standard Fourier shell correlation) criterion. Fourier shell correlations (FSC) between independently refined half-maps are shown before (blue) and after (red) post-processing; FSC between the map and the model is shown in black (Chen, McMullan et al., 2013).

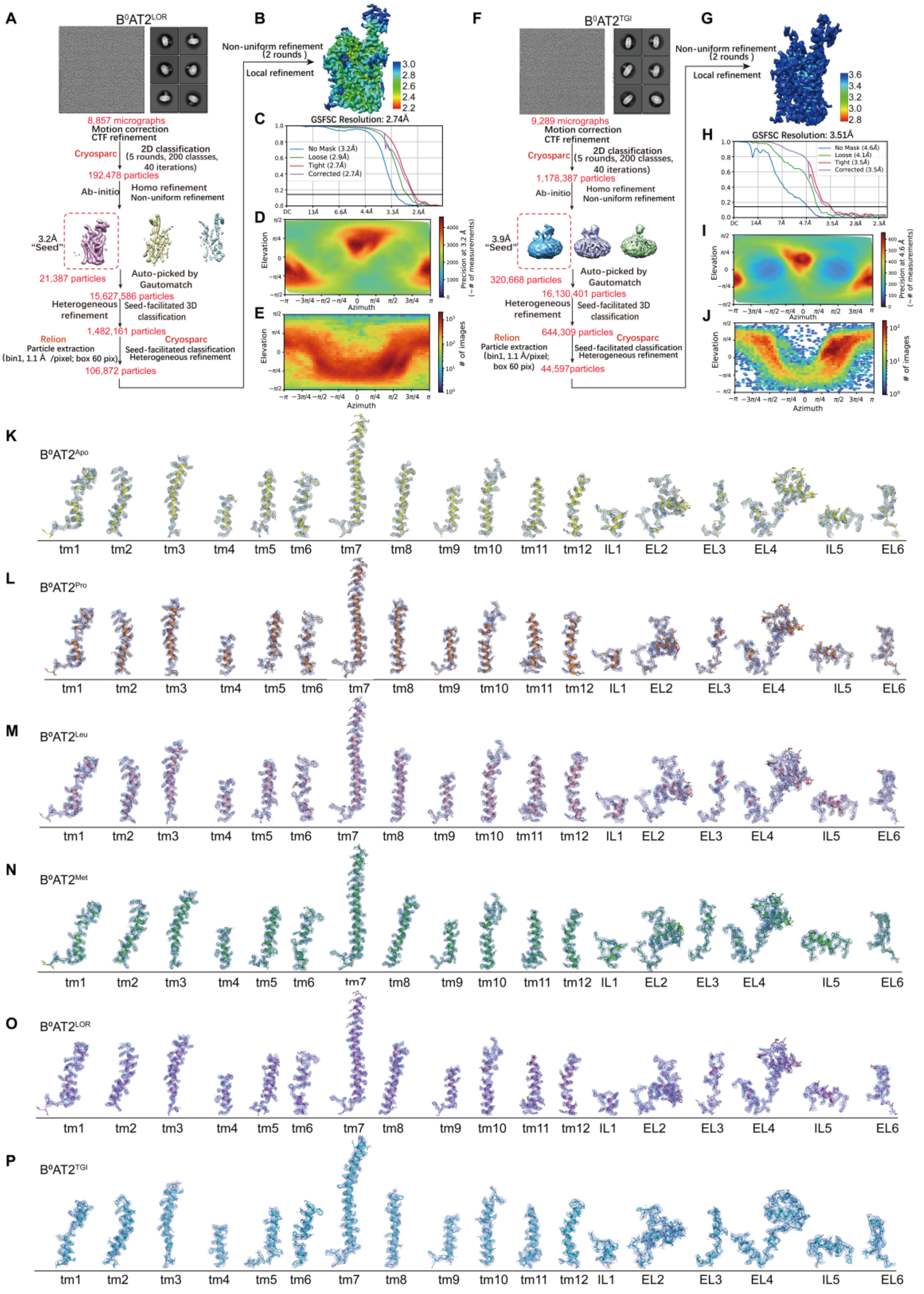

**Figure EV2. Cryo-EM data processing of $B^0AT2^{LOR}$ and $B^0AT2^{TGI}$ and the cryo-EM density and atomic model of TM1-12 of $B^0AT2$ in different states.**

(**A-J**) Flowchart for cryo-EM data processing of $B^0AT2^{LOR}$ and $B^0AT2^{TGI}$ with final cryo-EM maps colored by local resolution and angular distribution of the particles used in the final reconstruction. Representative raw cryo-EM micrograph and 2D class averages show distinct secondary structure features from different views of each structure. Several rounds of classifications are conducted to clean particles, followed by CTF refinement and local refinement to improve image quality. The final resolution is reported at 2.74 Å and 3.51 Å according to the GSFSC criterion. FSC of the final map for each structure are calculated between two independently refined half-maps before (blue) and after (red) post-processing. The FSC curve calculated between the cryo-EM density map and the corresponding structural model is shown in black.

(**K–P**) The cryo-EM density and atomic model of TM1-12, intracellular and extracellular loop of $B^0AT2^{Apo}$, $B^0AT2^{Pro}$, $B^0AT2^{Leu}$, $B^0AT2^{Met}$, $B^0AT2^{LOR}$ and $B^0AT2^{TGI}$.

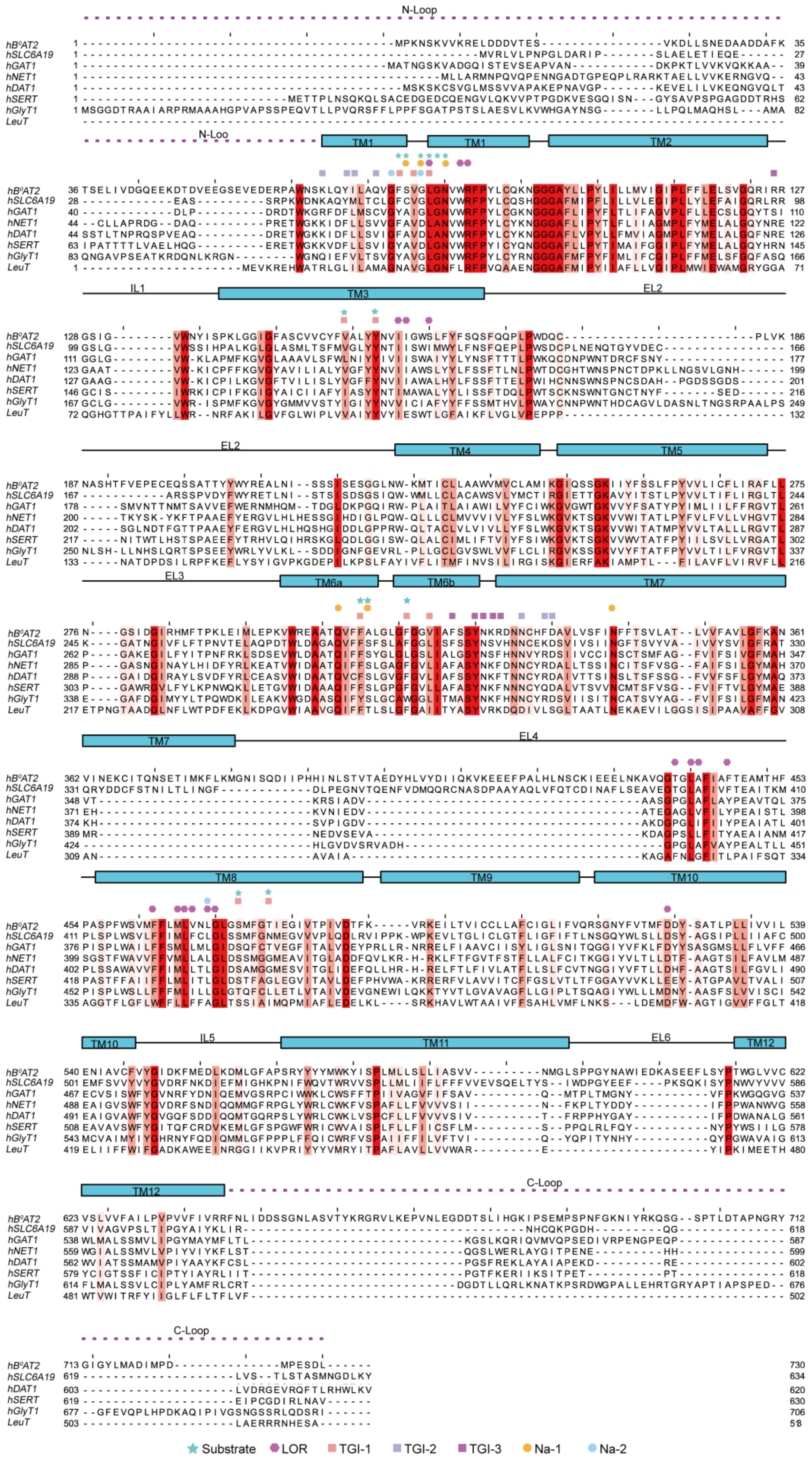
N-Loop
TM1
TM2
IL1
TM3
EL2
TM4
TM5
EL3
TM6a
TM6b
TM7
EL4
TM8
TM9
TM10
IL5
TM11
EL6
TM12
C-Loop
hB0AT2
hSLC6A19
hGAT1
hNET1
hDAT1
hSERT
hGlyT1
LeuT
Substrate
LOR
TGI-1
TGI-2
TGI-3
Na-1
Na-2

**Figure EV3. Sequence alignment of $B^0AT2$, SLC6A19, LeuT, and other representative NSS transporters.**

Secondary structural elements of $hB^0AT2$ (SLC6A15; UniProt: Q9H2J7) are depicted above the sequence alignment, and unmodeled loops are represented as dashed lines. Human Solute carrier family 6 member 19 (hSLC6A19, $B^0AT1$; UniProt: Q695T7), *Aquifex aeolicus* $Na^+$:neurotransmitter symporter (LeuT; UniProt: O67854), human dopamine transporter (hDAT, SLC6A3; UniProt: Q01959-1), human GABA transporter 1 (hGAT1, SLC6A1; UniProt: P30531-1), human serotonin transporter (hSERT, SLC6A4; UniProt: P31645-1), and human norepinephrine transporter (hNET, SLC6A2; UniProt: P23975-1) were aligned with $hB^0AT2$ using Clustal Omega and visualized with Jalview. Conserved residues among these proteins are highlighted in red or salmon, respectively. Residues involved in ligand binding are indicated by different symbols above the alignment.

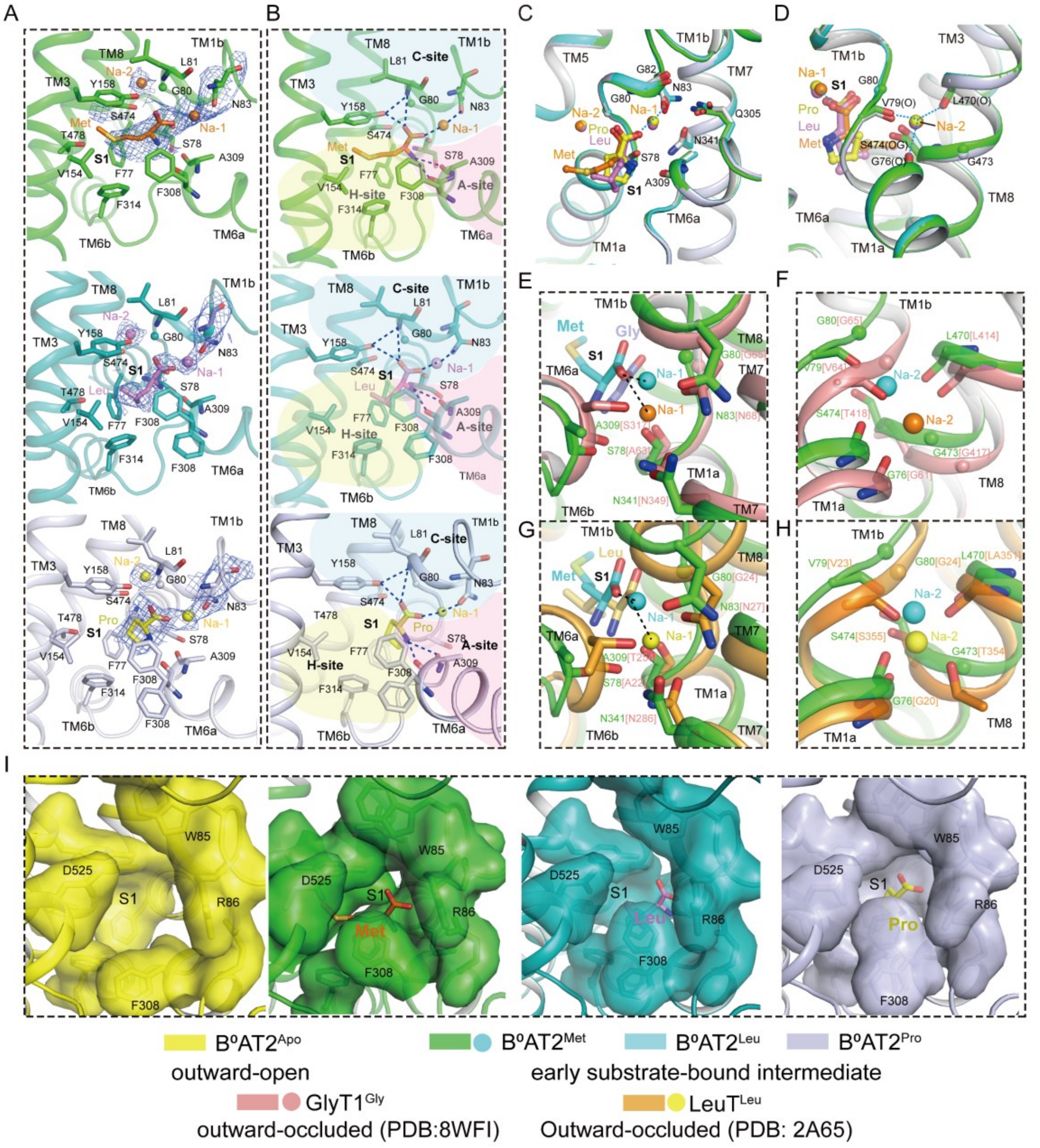

**Figure EV4. The substrate-binding site, ion-binding site, and extracellular channel above S1 controlled by the key gate residue Phe308.**

(**A**) Cryo-EM densities and binding modes of Met, Leu and Pro with associated modeled $Na^+$ ions. $Na^+$ ions are colored consistently with each substrate (orange: Met; violet: Leu; yellow: Pro). Cryo-EM densities are blue meshes.

(**B**) Substrate-binding pockets of $B^0$AT2 bound to Met, Leu, and Pro. Substrates are shown as sticks (orange, violet, yellow); modeled $Na^+$ ions as spheres. Interacting residues are shown as sticks colored by structure (green, cyan, blue-white). Hydrogen bonds and coordination bonds are shown as blue dashed lines. The amino group region, carboxylate region, and hydrophobic pocket are colored pink, teal, and pale yellow, respectively.

(**C–D**) Superposition of structures with modeled Na-1 (C) and modeled Na-2 (D). Substrates and interacting residues are shown as sticks (Leu, violet; Met, green; Pro, yellow); modeled $Na^+$ ions and are shown in matching colors.

(**E-H**) Superposition of the Na-1 (E, G) and Na-2 (F, H) binding sites modeled in $B^0AT2^{Met}$ (green), $GlyT1^{Gly}$ (salmon, PDB 8WFI) (Wei, Li et al., 2024), and $LeuT^{Leu}$ (orange, PDB 2A65) (Yamashita, Singh et al., 2005). $Na^+$ ions are shown as spheres (cyan, modeled in $B^0AT2$; orange, GlyT1; yellow, LeuT). Substrates are shown as sticks (Met, cyan; Gly, light blue; Leu, yellow). The residues mediating the interactions are indicated, with corresponding residues at the same positions shown in square brackets, and coordination bonds depicted as dashed lines.

(**I**) Comparison of the extracellular channel above the S1 pocket in $B^0AT2^{Apo}$ and $B^0AT2^{Met}$, $B^0AT2^{Leu}$, $B^0AT2^{Pro}$ complexes, controlled by the key gate residue Phe308. The conformational states are outward-open in the apo form, early substrate-bound intermediate in $B^0AT2^{Met}$, $B^0AT2^{Leu}$, and $B^0AT2^{Pro}$.

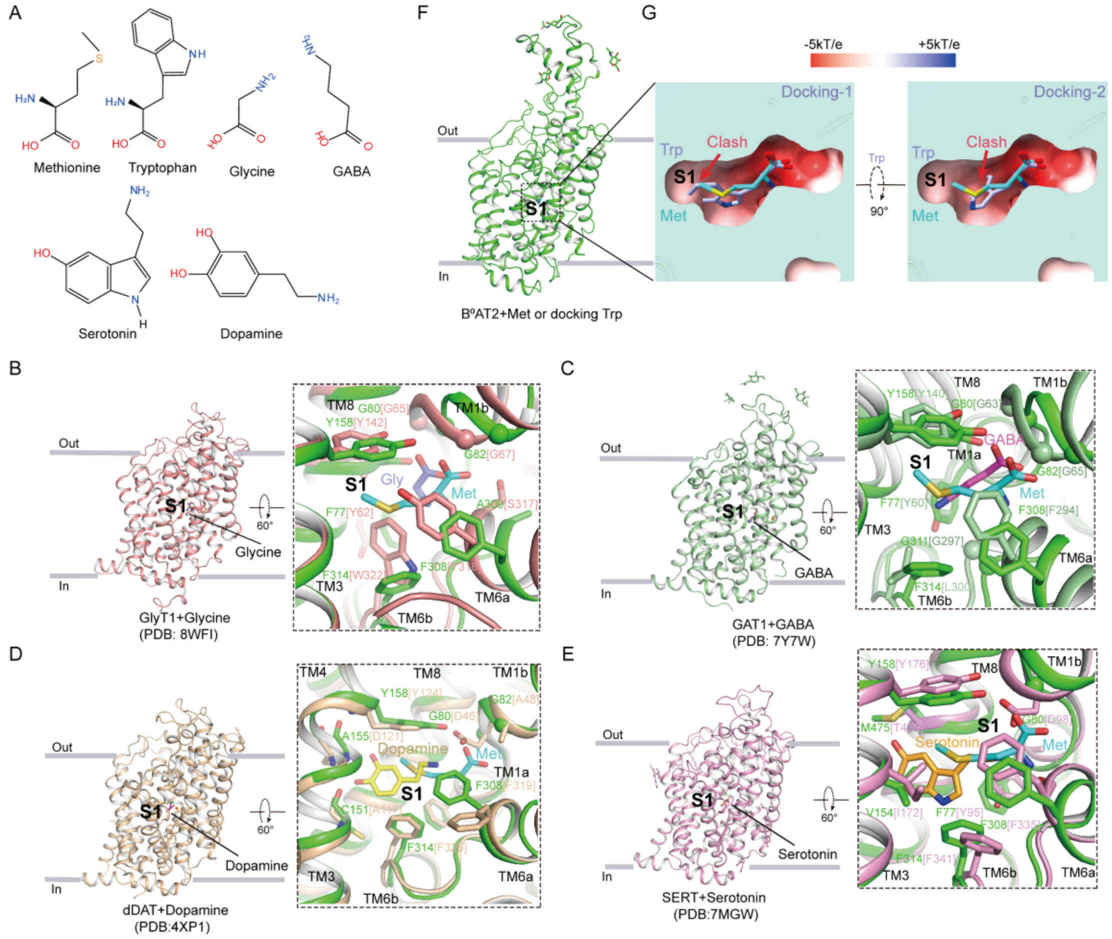

**Figure EV5. Substrate selectivity and structural divergence of S1 pockets in B$^0$AT2 and representative NSS transporters.**

(**A**) Chemical structures of Met, Trp, Gly, GABA, serotonin, and dopamine.

(**B-C**) Comparison of the amino acid transporter B$^0$AT2$^{Met}$ with Gly-bound GlyT1 (PDB 8WFI) (Wei et al., 2024) and GABA-bound GAT1 (PDB 7Y7W) (Zhu, Huang et al., 2023). Conserved residues Gly80 and Gly82 are shown as spheres, forming a spacious cavity to accommodate the carboxylate head group of amino acid substrates.

(**D-E**) Comparison of B$^0$AT2$^{Met}$ with monoamine transporters dopamine-bound dDAT (PDB 4XP1) (Wang, Penmatsa et al., 2015) and serotonin-bound SERT (PDB 7MGW) (Gouaux, 2021). Gly80 is replaced by Asp46 (dDAT) and Asp98 (SERT), resulting in steric clashes that prevent amino acid substrate binding.

(**F-G**) Comparison of Met-bound B$^0$AT2 and Trp-docked B$^0$AT2 model. Electrostatic surface representation showing that docking Trp into the Met-bound S1 pocket results in severe steric clashes, even after rotation into different orientations.

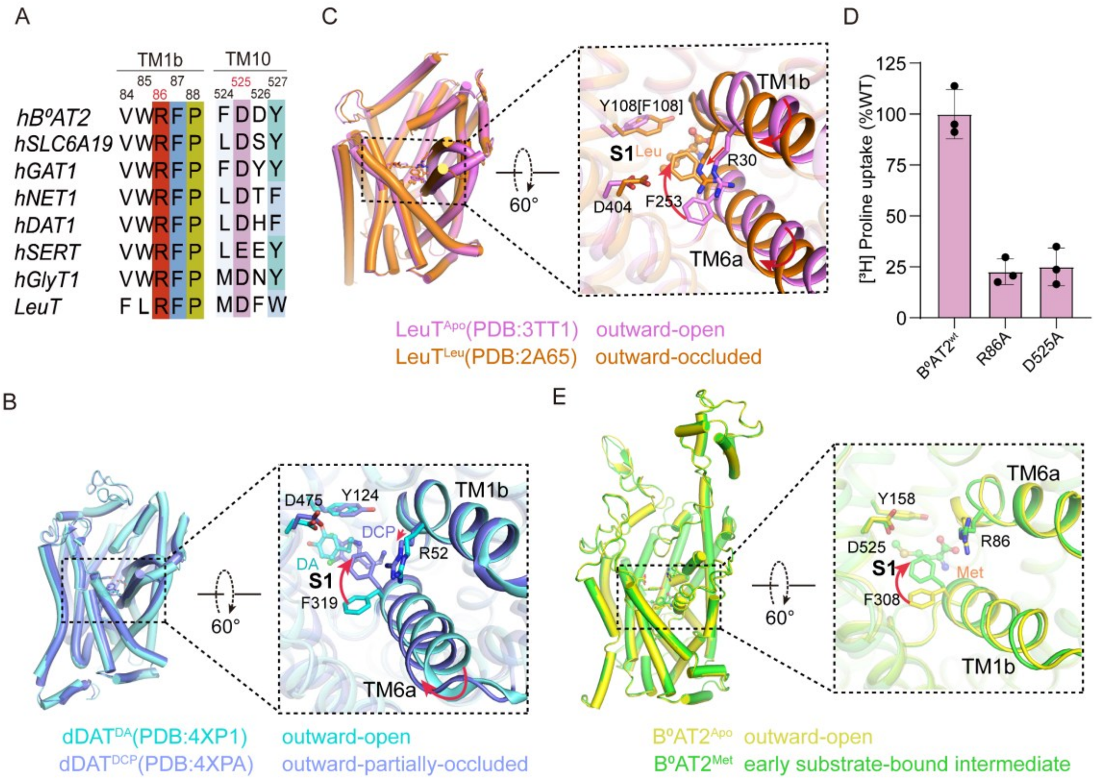

**Figure EV6. Substrate binding induces structural rearrangements in the S1 pocket.**

(**A**) Sequence alignment of B$^0$AT2 with representative NSS members, SLC6A19 and LeuT shows that the extracellular gate residues Arg86 and Asp525 are highly conserved. (**B**) Superposition of dopamine-bound dDAT (dDAT$^{DA}$; PDB 4XP1) (Wang et al., 2015) in the outward-open conformation and DCP-bound dDAT (dDAT$^{DCP}$; PDB 4XPA) (Wang et al., 2015) in the outward-partially occluded conformation, shown in cyan and slate, respectively. Black dashed boxes highlight TM1b and TM6a of the extracellular vestibule. Key residues and ligands are shown as sticks, with structural features shown in the right dashed box. Red arrows indicate the direction of pronounced displacement. (**C**) Superposition of the outward-open state of LeuT in apo state (LeuT$^{Apo}$; PDB 3TT1) (Harini Krishnamurthy1 & Eric Gouaux1, Krishnamurthy et al., 2012) and the outward-occluded state of Leu-bound LeuT (LeuT$^{Leu}$; PDB 2A65) (Yamashita et al., 2005), shown in violet and orange, respectively. Black dashed boxes highlight TM1b and TM6a of the extracellular vestibule. Key residues and ligands are shown as sticks, with structural features shown in the right dashed box. Red arrows indicate the direction of pronounced displacement.

(**D**) Transport activity of WT $B^0AT2$ and mutants R86A and D525A quantified using [3H]-proline uptake assay. The transport activity of each mutant is normalized to that of WT $B^0AT2$. Data represent mean ± s.d. (error bars), n = 3 technical replicates. The experiment was performed independently twice with similar results.

(**E**) Superposition of the outward-open state of $B^0AT2^{Apo}$and the early substrate-bound intermediate state of $B^0AT2^{Met}$, shown in yellow and green, respectively. Black dashed boxes highlight TM1b and TM6a of the extracellular vestibule. Key residues and ligands are shown as sticks, with structural features shown in the right dashed box. Red arrows indicate the direction of pronounced displacement.

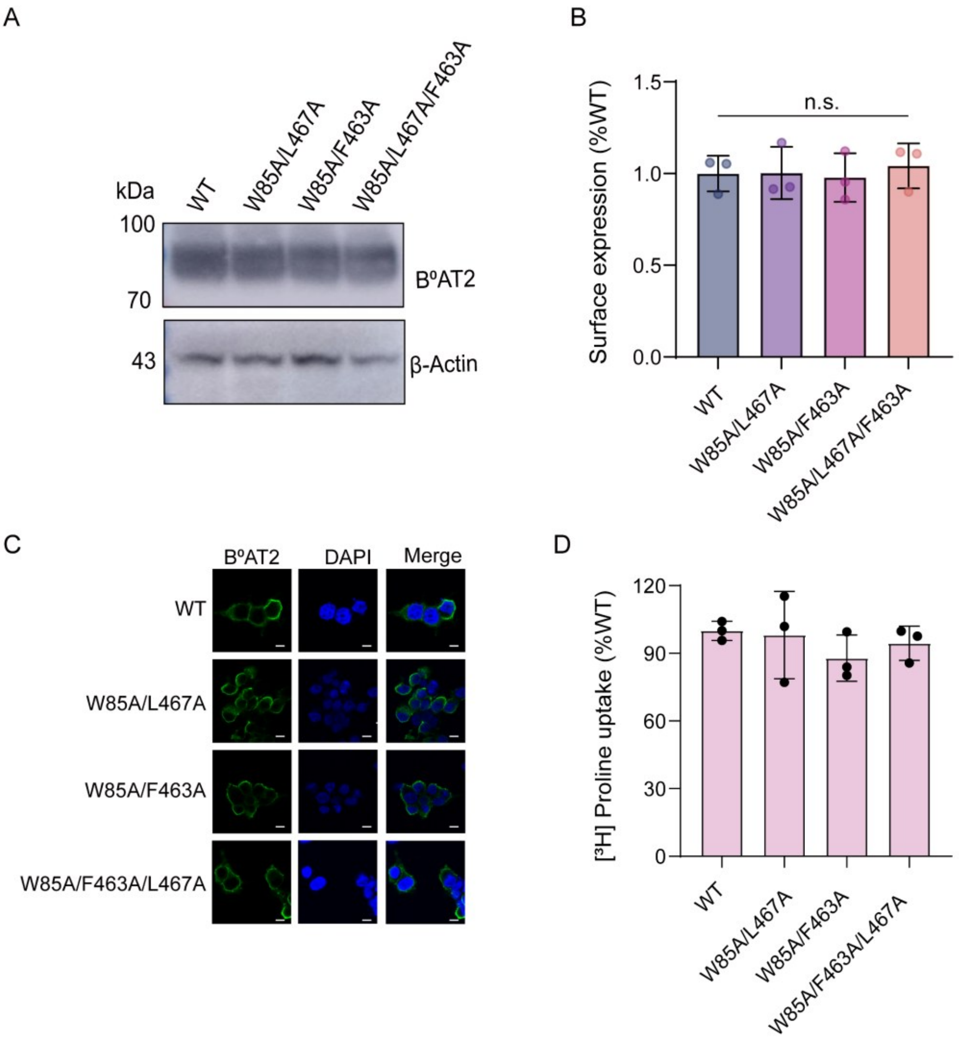

**Figure EV7. Expression levels, localization, and transport activity analyzed by immunostaining, western blotting and transport assay expressing WT B$^0$AT2 or mutants.**

(**A**) Representative Western blot images showing the expression of WT and mutant proteins. β-Actin is used as a loading control. The experiment was performed independently twice with similar results.

(**B**) Quantification of Western blot band intensities, normalized to β-Actin, showing relative expression levels of each mutant compared to WT B$^0$AT2. Each point represents an independent experiment. Statistical analysis was performed using one-way ANOVA with appropriate post hoc tests. n.s., not significant ($P \geq 0.05$).

(**C**). Representative images of anti-FLAG immunostaining in B$^0$AT2 and mutants transfected HEK-293T cells. Scale bar = 5 μm.

(**D**). Transport activity of WT $B^0AT2$ and mutants quantified using [$^3$H]-proline uptake assay. The transport activity of each mutant is normalized to that of WT $B^0AT2$. The uptake duration is 1 min to ensure the uptake occurs within the linear range. Data represent mean ± s.d. (error bars), n = 3 technical replicates. The experiment was performed independently twice with similar results.

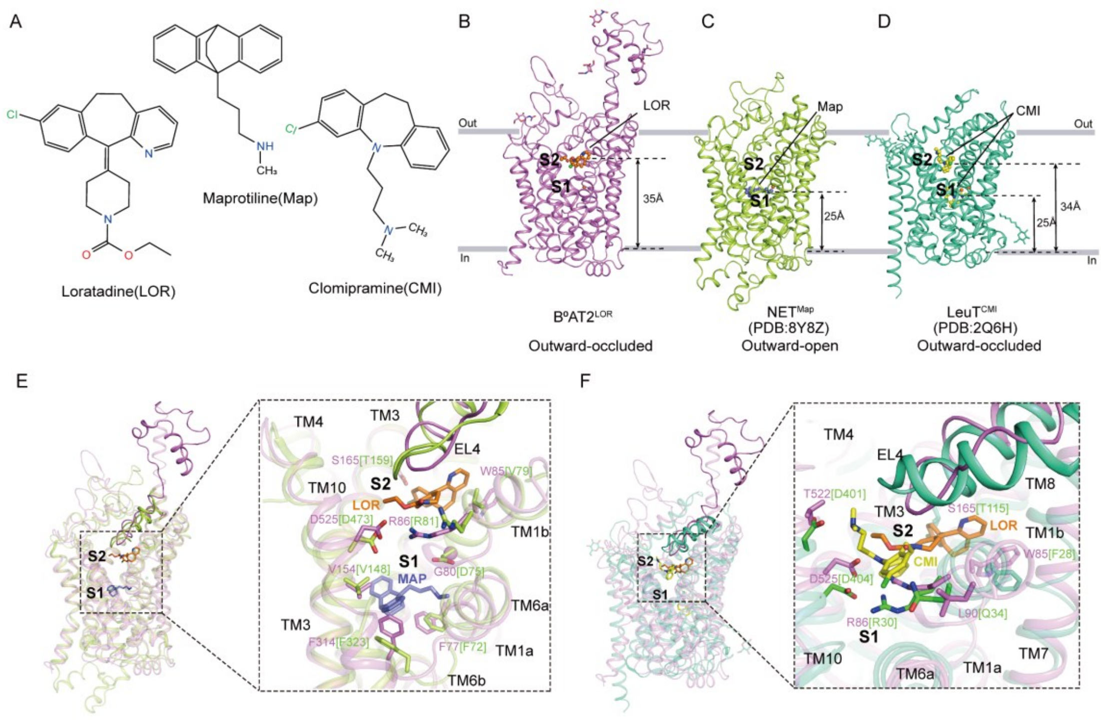

**Figure EV8. Comparison of binding sites of LOR, Map, and CMI in B$^0$AT2, NET and LeuT.**

(**A**) Chemical structures of loratadine (LOR), maprotiline (Map), and clomipramine (CMI).

(**B–D**) Overall structures of B$^0$AT2$^{LOR}$ (violet), NET$^{Map}$ (PDB 8Y8Z, lime) (Zhang, Yin et al., 2024), and LeuT$^{CMI}$ (PDB 2Q6H, green-cyan) (Singh, Yamashita et al., 2007) in the outward-occluded, outward-open, and outward-occluded conformational states, respectively, shown in cartoon representation. Inhibitors are shown as sticks: LOR (orange), Map (light-blue), and CMI (yellow). Distances from the extracellular surface to key residues are indicated, and conformational states are annotated.

(**E**) Comparison of the LOR- and Map-binding pockets in B$^0$AT2 and NET**.** Key interacting residues are shown as sticks and labeled. LOR and Map are depicted as orange and light-blue sticks, respectively. Residues involved in gating and substrate coordination (TM1b, TM3, TM6a, TM10, and EL4) are highlighted to illustrate differences in inhibitor-binding environments and conformational adaptations.

(**F**) Comparison of the LOR- and CMI-binding pockets in B$^0$AT2 and LeuT. Key interacting residues are shown as sticks and labeled. LOR and CMI are represented as orange and yellow sticks, respectively. Structural features of the binding pockets highlight conserved and divergent interactions underlying inhibitor recognition and specificity.

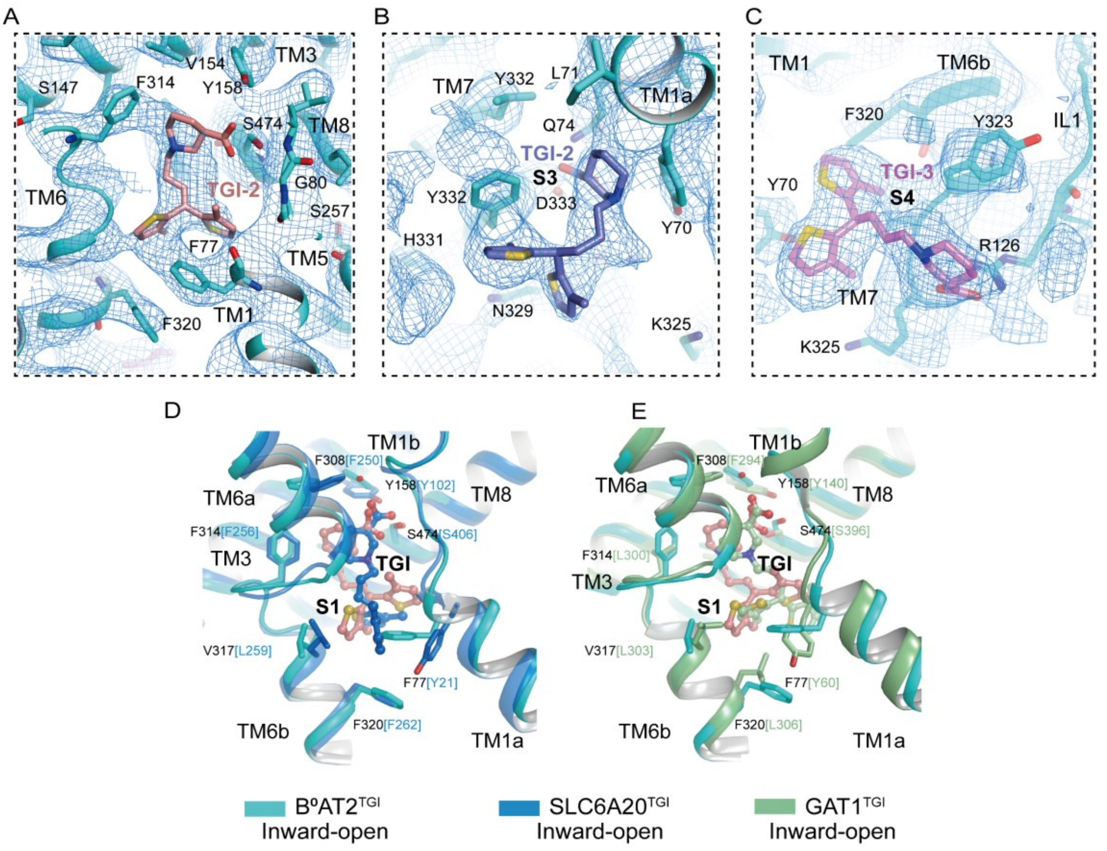

**Figure EV9. Structural comparison of B$^0$AT2, SLC6A20 and GAT1 bound with TGI.**

(A-C) Cryo-EM density maps (blue mesh) and atomic models of TGI-1 (salmon sticks), TGI-2 (slate sticks), and TGI-3 (violet sticks) bound within the S1 (A), S3 (B), and S4 (C) pockets in B$^0$AT2$^{TGI}$, respectively.

(**D**). Structural alignment of inward-open B$^0$AT2$^{TGI}$ (cyan) and SLC6A20$^{TGI}$ (marine, PDB 8WM3) (Broer, Hu et al., 2024). TGI in B$^0$AT2 and in SLC6A20 are shown as sticks (salmon, marine). S1 site residues are displayed as sticks and colored by protein.

(**E**). Structural alignment of inward-open B$^0$AT2$^{TGI}$ (cyan) and GAT1$^{TGI}$ (pale green, PDB 7Y7Z) (Zhu et al., 2023). TGI in B$^0$AT2 and in GAT1 are shown as sticks (salmon, marine). S1 site residues are displayed as sticks and colored by protein.

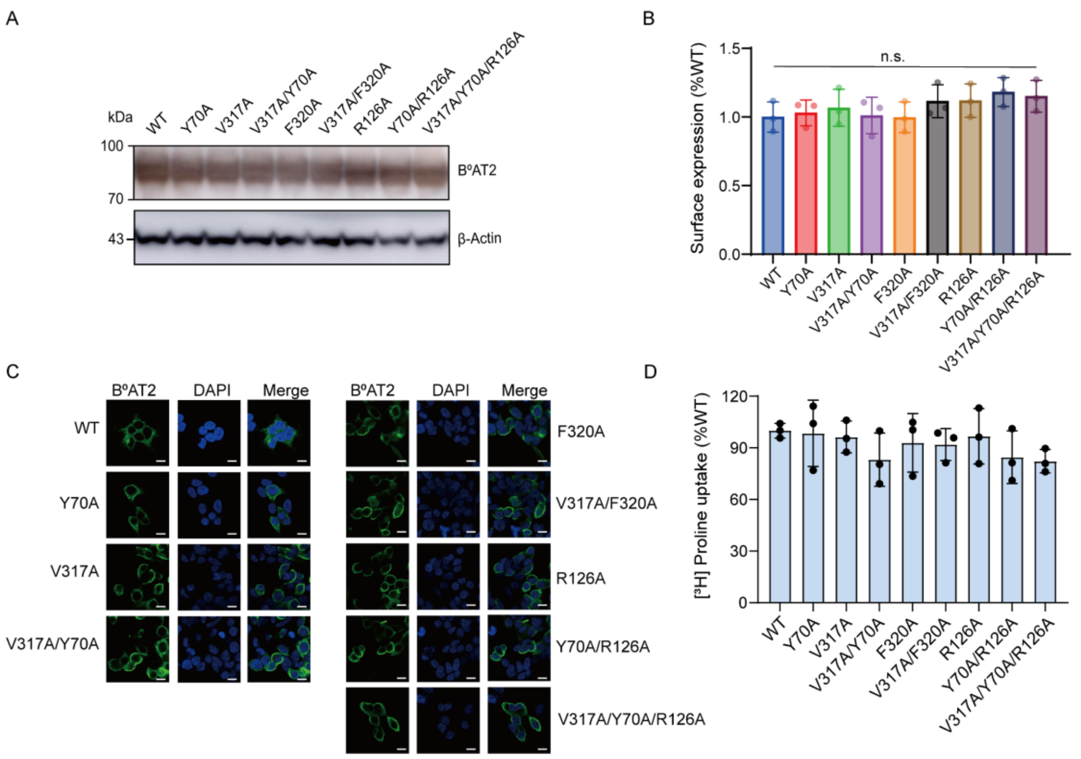

**Figure EV10. Expression levels, localization, and transport activity analyzed by immunostaining, western blotting and transport assay expressing WT B$^0$AT2 or mutants.**

(**A**) Representative Western blot images showing the expression of WT and mutant proteins. β-Actin is used as a loading control. The experiment was performed independently twice with similar results.

(**B**) Quantification of Western blot band intensities, normalized to β-Actin, showing relative expression levels of each mutant compared to WT B$^0$AT2. Each point represents an independent experiment. Statistical analysis was performed using one-way ANOVA with appropriate post hoc tests. n.s., not significant ($P \geq 0.05$).

(**C**). Representative images of anti-FLAG immunostaining in B$^0$AT2 and mutants transfected HEK-293T cells. Scale bar = 5 μm.

(**D**). Transport activity of WT B$^0$AT2 and mutants quantified using [$^3$H]-proline uptake assay. The transport activity of each mutant is normalized to that of WT B$^0$AT2. The uptake duration is 1 min to ensure the uptake occurs within the linear range. Data represent mean ± s.d. (error bars), n = 3 technical replicates. The experiment was performed independently twice with similar results.

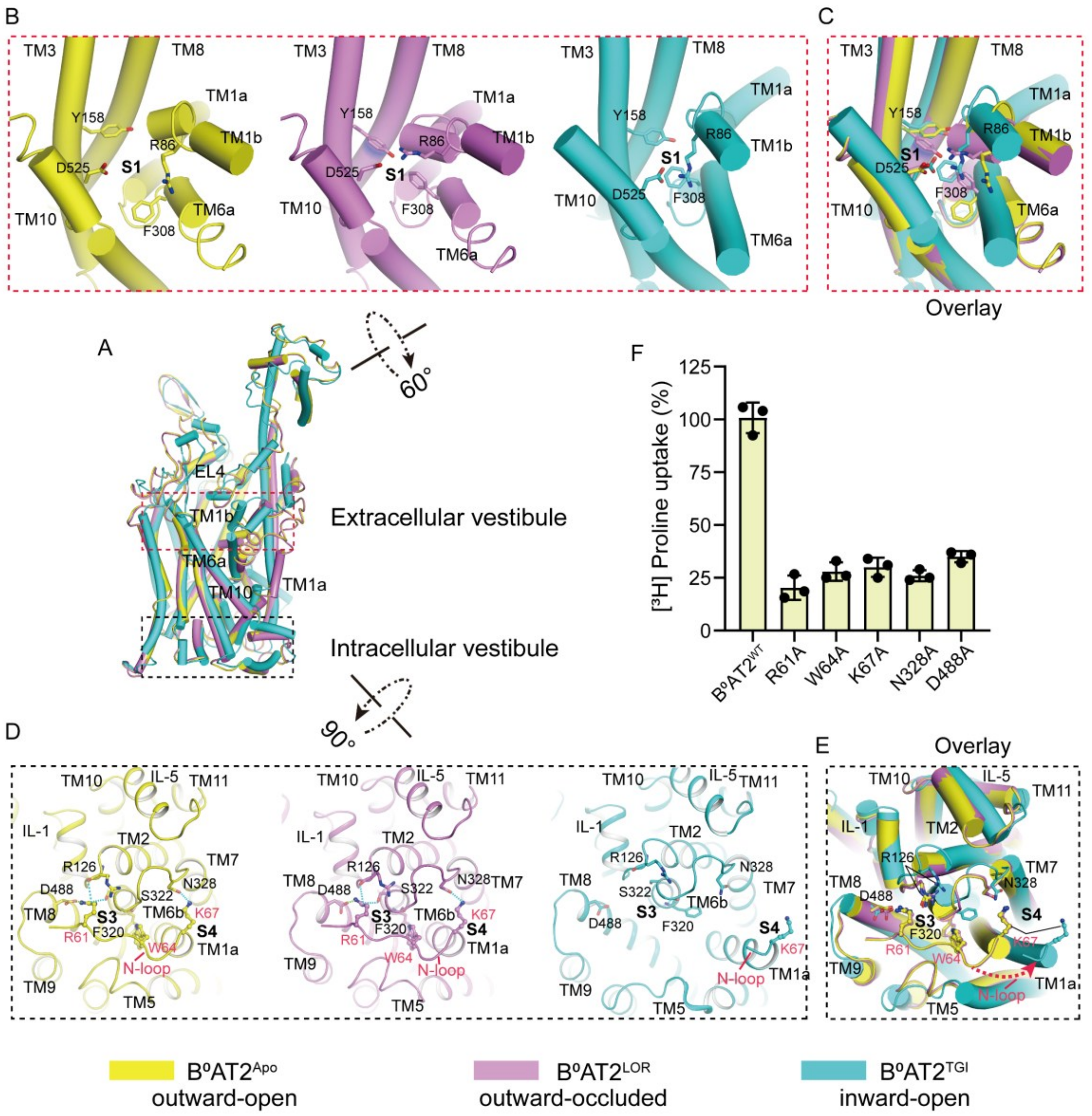

**Figure EV11. Interactions stabilizing distinct conformational states and gating regulation in B$^0$AT2.**

(**A**) Structural superposition of B$^0$AT2$^{Apo}$ (yellow), B$^0$AT2$^{LOR}$ (violet) and B$^0$AT2$^{TGI}$ (cyan), illustrating distinct global conformational states of the transporter. Red and black dashed boxes denote the extracellular vestibule and intracellular vestibule, respectively, whose structural features are compared in panels (B)–(E).

(**B**) Close-up views of the extracellular vestibule above the S1 pocket in the three conformational states. In the B$^0$AT2$^{Apo}$ structure, the extracellular vestibule remains open, consistent with an outward-open conformation. In B$^0$AT2$^{LOR}$, the extracellular vestibule is sealed by a conserved gating network involving Arg86, Asp525, Phe308, and Tyr158, defining an outward-occluded state. In B$^0$AT2$^{TGI}$, the same residues maintain closure of the extracellular vestibule, while the transporter adopts an inward-open conformation.

(**C**) Superimposed extracellular views of the three structures show that TM1b, TM6a and TM10 move further toward the center of the transport pathway in $B^0AT2^{TGI}$ compared with $B^0AT2^{Apo}$ and $B^0AT2^{LOR}$, thereby further stabilizing closure of the extracellular vestibule.

(**D**) Comparison of the intracellular vestibule architecture reveals a critical role for the N-loop (residues 59–64) in stabilizing cytoplasmic closure. In $B^0AT2^{Apo}$ and $B^0AT2^{LOR}$, N-loop residues Arg61, Trp64, and Lys67 form extensive interactions with intracellular helices, resulting in a closed intracellular vestibule. In $B^0AT2^{TGI}$, elevation of TM1a disrupts these interactions, leading to the loss of N-loop density and opening of the intracellular vestibule.

(**E**) Superposition of the intracellular vestibule shown in cylinder representation, with key residues displayed as sticks. The red dashed arrow indicates the direction of N-loop displacement associated with transition to the inward-open conformation.

(**F**) Functional validation of intracellular gate residues. [$^3$H]-proline uptake assays show that mutations of N-loop and intracellular gate residues markedly reduce transport activity, underscoring their essential role in intracellular vestibule integrity and conformational coupling. Data represent mean ± s.d. (error bars), n = 3 technical replicates. The experiment was performed independently twice with similar results.

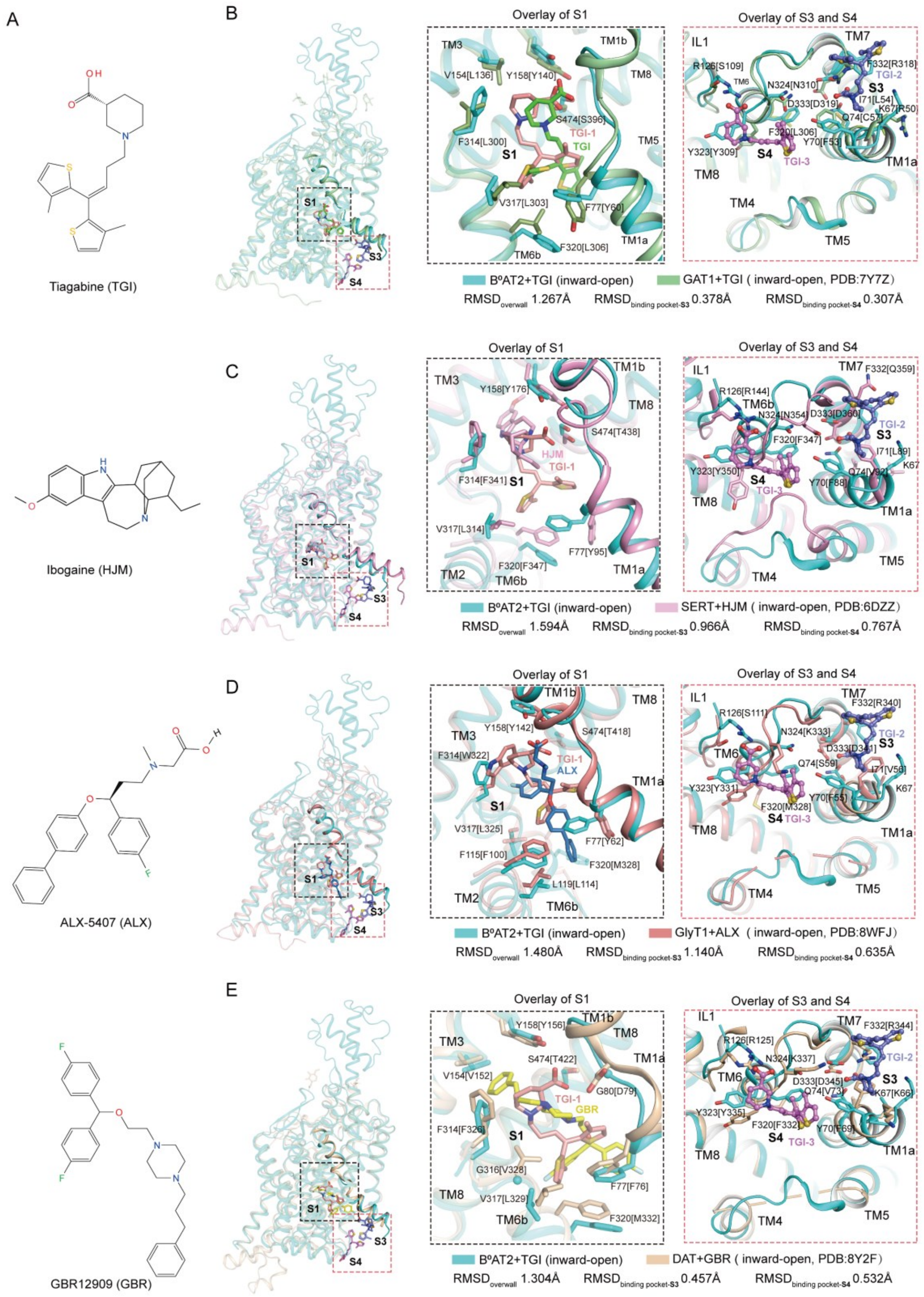

**Figure EV12. Structural comparison of S1, S3 and S4 in the inward-open state of $B^0AT2^{TGI}$ and other representative NSS transporters.**

(**A**) The chemical structures of TGI, HJM, ALX-5407, and GBR12909 are depicted.

(**B**) Structural alignment of inward-open $B^0AT2^{TGI}$ (cyan) and $GAT1^{TGI}$ (pale green,

PDB 7Y7Z) (Zhu et al., 2023). TGI-1–3 in $B^0AT2$ and TGI in GAT1 are shown as sticks (salmon, slate, violet, and green). S1, S3, and S4 site residues are displayed as sticks and colored by protein, with enlarged views provided. The overall RMSD is 1.267 Å, with S3 and S4 pocket RMSDs of 0.378 Å and 0.307 Å, respectively.

(**C**) Structural alignment of inward-open $B^0AT2^{TGI}$ (cyan) and $SERT^{HJM}$ (pink, PDB 6DZZ) (Coleman, Yang et al., 2019). TGI-1–3 in $B^0AT2$ and HJM in SERT are shown as sticks (salmon, slate, violet, and pink). S1–S4 site residues are displayed as sticks and colored by protein, with enlarged views provided. The overall RMSD is 1.594 Å, with S3 and S4 pocket RMSDs of 0.966 Å and 0.767 Å, respectively.

(**D**) Structural alignment of inward-open $B^0AT2^{TGI}$ (cyan) and $GlyT1^{ALX}$ (salmon, PDB 8WFJ) (Wei et al., 2024). TGI-1–3 in $B^0AT2$ and ALX in GlyT1 are shown as sticks (salmon, slate, violet, and sky-blue). S1, S3, and S4 site residues are displayed as sticks and colored by protein, with enlarged views provided. The overall RMSD is 1.480 Å, with S3 and S4 pocket RMSDs of 1.140 Å and 0.635 Å, respectively.

(**E**) Structural alignment of inward-open $B^0AT2^{TGI}$ (cyan) and $DAT^{GBR}$ (wheat, PDB 8Y2F) (Li, Wang et al., 2024). TGI-1–3 in $B^0AT2$ and GBR in DAT are shown as sticks (salmon, slate, violet, and yellow). S1, S3, and S4 site residues are displayed as sticks and colored by protein, with enlarged views provided. The overall RMSD is 1.304 Å, with S3 and S4 pocket RMSDs of 0.457 Å and 0.532 Å, respectively.

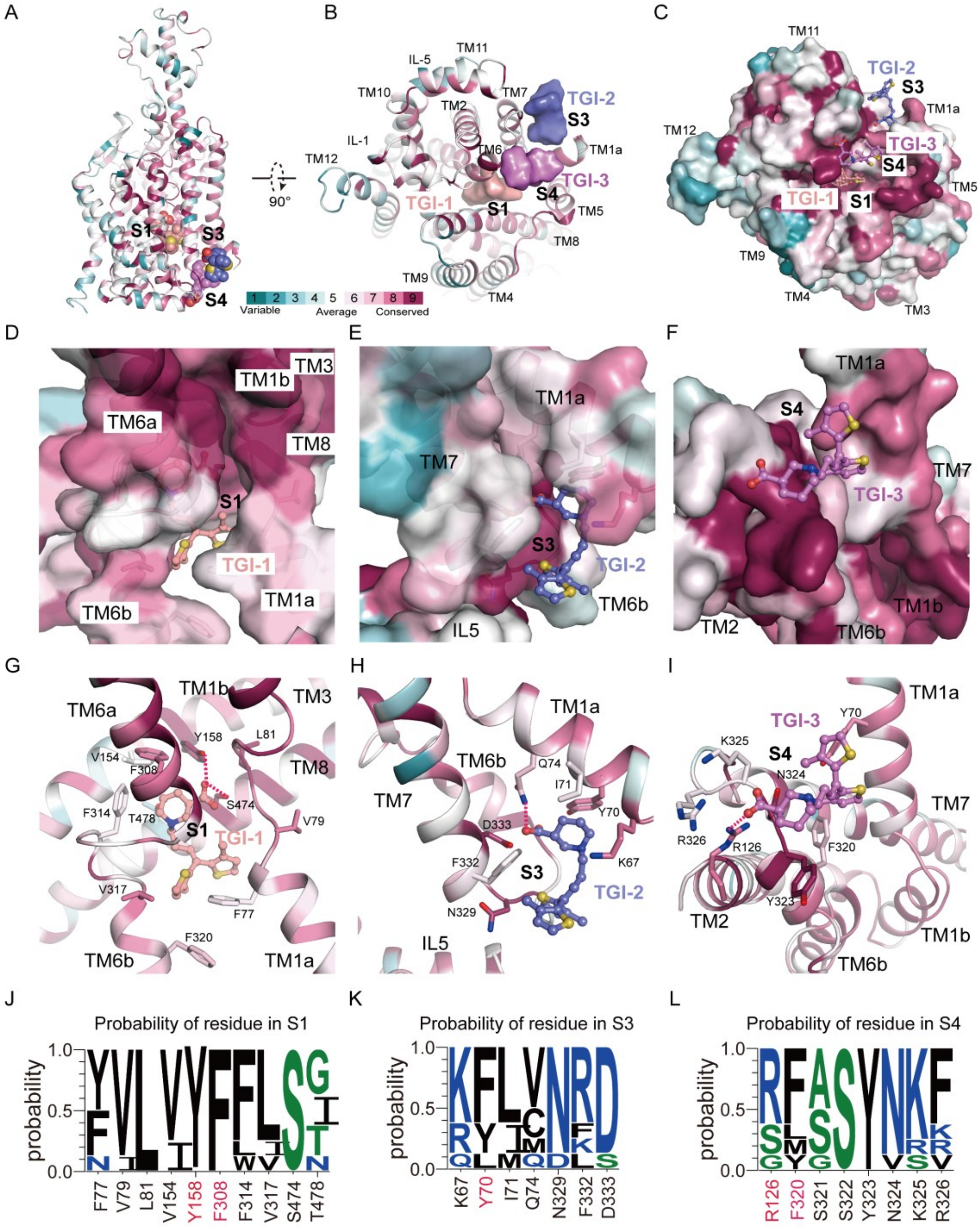

**Figure EV13. Sequence conservation analysis of B⁰AT2 across the SLC6 family.**

(**A–B**) Overall view of sequence conservation mapped onto the human $B^0AT2$ structure in complex with TGI. The protein is shown in cartoon representation, whereas the three TGI molecules are rendered as surfaces to highlight their binding sites. Two orthogonal views are presented to illustrate the global distribution of conserved residues. Sequence conservation among SLC6 family members (excluding SLC6A11) was calculated using

ConSurf (Ashkenazy, Abadi et al., 2016) and mapped onto the protein surface, with red and cyan indicating high and low conservation, respectively.

(**C**) Surface view showing the highly conserved substrate-binding pockets S1, S3, and S4. TGI-1, TGI-2, and TGI-3 are shown in ball-and-stick representation.

(**D–F**) Close-up views of the S1, S3, and S4 pockets. The transporter is shown in cartoon and surface representations with sequence conservation mapped onto the surface. TGI-1, TGI-2, and TGI-3 are shown as ball-and-stick models in salmon, slate, and violet, respectively. Key transmembrane helices contributing to each pocket are labeled.

(**G–I**) Close-up views of the S1, S3, and S4 pockets highlighting interacting residues. The transporter is shown in cartoon representation with sequence conservation mapped onto the surface. Key residues forming each pocket are labeled, and the views correspond to those shown in (D–F).

(**J–L**) WebLogo (Crooks, Hon et al., 2004) representations showing residue probabilities at positions within the S1, S3, and S4 pockets. Letter height reflects amino acid frequency across $B^0$AT2, SLC6A19, LeuT, and representative NSS members (GAT1, NET, DAT, SERT, and GlyT1), consistent with structural conservation patterns. Residues highlighted in red denote representative residues.

#### Tables

**Table EV1. Cryo-EM data collection, refinement and validation statistics**

| | $B^0AT2^{Apo}$ (EMDB-68684) (PDB 22UJ) | $B^0AT2^{Pro}$ (EMDB-68745) (PDB 22XA) | $B^0AT2^{Leu}$ (EMDB-68740) (PDB 22WU) | $B^0AT2^{Met}$ (EMDB-68705) (PDB 22VB) | $B^0AT2^{LOR}$ (EMDB-68730) (PDB 22WA) | $B^0AT2^{TGI}$ (EMDB-68754) (PDB 22XL) |
|---|---|---|---|---|---|---|
| Magnification | 81,000X | 81,000X | 81,000X | 81,000X | 81,000X | 81,000X |
| Voltage (kV) | 300 | 300 | 300 | 300 | 300 | 300 |
| Electron exposure ($e^-/Å^2$) | 50 | 50 | 50 | 50 | 50 | 50 |
| Defocus range (μm) | -1.2~-1.8 | -1.2~-1.8 | -1.2~-1.8 | -1.2~-1.8 | -1.2~-1.8 | -1.2~-1.8 |
| Pixel size (Å) | 0.55 | 0.55 | 0.55 | 0.55 | 0.55 | 0.55 |
| Symmetry imposed | C1 | C1 | C1 | C1 | C1 | C1 |
| Initial particle images (no.) | 10,329,738 | 11,115,705 | 13,694,783 | 16,447,141 | 15,627,586 | 16,130,401 |
| Final particle images (no.) | 186,351 | 537,053 | 475,398 | 385,386 | 106,872 | 44,597 |
| Map resolution (Å) | 2.84 | 2.80 | 2.82 | 2.92 | 2.74 | 3.51 |
| FSC threshold | 0.143 | 0.143 | 0.143 | 0.143 | 0.143 | 0.143 |
| Map resolution range (Å) | 2.8-3.3 | 1.1-3.0 | 2.8-3.0 | 2.9-3.0 | 1.3-3.2 | 3.5-4.1 |
| Refinement | | | | | | |
| Initial model used | AlphaFold | $B^0AT2^{Apo}$ | $B^0AT2^{Apo}$ | $B^0AT2^{Apo}$ | $B^0AT2^{Apo}$ | De novo build |
| Model resolution (Å) | 2.84 | 2.80 | 2.82 | 2.92 | 2.74 | 3.51 |
| FSC threshold | 0.5 | 0.5 | 0.5 | 0.5 | 0.5 | 0.5 |
| Map sharpening B factor (Å2) | 131.8 | 117.4 | 115.9 | 111.1 | 129.4 | 102.1 |
| Model composition | | | | | | |
| Non-hydrogen atoms | 4,741 | 4,760 | 4,776 | 4,763 | 4,768 | 4,784 |
| Protein residues | 589 | 589 | 589 | 589 | 589 | 582 |
| Ligands | 4 | 5 | 5 | 5 | 5 | 4 |

| Table EV1. Cryo-EM data collection, refinement and validation statistics (continued) | | | | | | |
|---|---|---|---|---|---|---|
| B factors (Å$^2$) | | | | | | |
| Protein | 62.93 | 37.79 | 95.46 | 110.07 | 39.72 | 110.71 |
| Ligand | 116.80 | 83.35 | 144.26 | 158.54 | 68.11 | 111.56 |
| R.m.s. deviations | | | | | | |
| Bond lengths (Å) | 0.003 | 0.002 | 0.002 | 0.003 | 0.003 | 0.003 |
| Bond angles (°) | 0.540 | 0.436 | 0.451 | 0.470 | 0.568 | 0.662 |
| Validation | | | | | | |
| MolProbity score | 2.10 | 2.18 | 2.05 | 2.37 | 1.58 | 2.52 |
| Clashscore | 4.63 | 15.10 | 12.26 | 13.65 | 3.77 | 39.16 |
| Poor rotamers (%) | 5.24 | 1.74 | 1.55 | 3.69 | 2.14 | 0.39 |
| Ramachandran plot | | | | | | |
| Favored (%) | 95.40 | 95.57 | 95.57 | 95.74 | 97.10 | 93.10 |
| Allowed (%) | 4.60 | 4.43 | 4.43 | 4.26 | 2.90 | 6.55 |
| Disallowed (%) | 0.00 | 0.00 | 0.00 | 0.00 | 0.00 | 0.34 |

#### Legends for Movies EV1 to EV3

**Movie EV1 (separate file). Structural comparison of apo and Met-bound $B^0$AT2 revealing early substrate recognition.** This movie shows $B^0$AT2 transitioning from the apo outward-open state to the Met-bound early substrate-bound intermediate. Upon Met binding, the key gating residue Phe308 undergoes rotameric rearrangement, closing the extracellular entrance to S1 and remodeling pocket geometry to accommodate substrate. This local rearrangement stabilizes substrate binding and captures a transient early intermediate during substrate recognition.

**Movie EV2 (separate file). Structural comparison of apo and LOR-bound $B^0$AT2 revealing allosteric stabilization of the outward-occluded state.** This movie superimposes the apo outward-open and LOR-bound outward-occluded states of $B^0$AT2. LOR binding at the extracellular S2 allosteric pocket remodels the S1–S2 gating network, blocks the extracellular entrance to the S1 pocket, and stabilizes the outward-occluded conformation to mediate allosteric inhibition.

**Movie EV3 (separate file). Movie S3. Structural comparison of apo and TGI-bound $B^0$AT2 reveals cooperative multi-site inhibition via intracellular cavity engagement.** This movie compares the apo outward-open and TGI-bound inward-open states of $B^0$AT2. TGI-1, TGI-2, and TGI-3 occupy the S1, S3, and S4 pockets, remodeling the intracellular vestibule and rearranging key gating residues. Intracellularly, TGI binding drives TM1a displacement, prevents its conformational reset, and locks $B^0$AT2 in an inhibited inward-open state through cooperative multi-site engagement.

#### Supplementary References

Ashkenazy H, Abadi S, Martz E, Chay O, Mayrose I, Pupko T, Ben-Tal N (2016) ConSurf 2016: an improved methodology to estimate and visualize evolutionary conservation in macromolecules. *Nucleic Acids Research* 44: W344-W350

Broer A, Hu Z, Kukulowicz J, Yadav A, Zhang T, Dai L, Bajda M, Yan R, Broer S (2024) Cryo-EM structure of ACE2-SIT1 in complex with tiagabine. *J Biol Chem* 300: 107687

Chen S, McMullan G, Faruqi AR, Murshudov GN, Short JM, Scheres SH, Henderson R (2013) High-resolution noise substitution to measure overfitting and validate resolution in 3D structure determination by single particle electron cryomicroscopy. *Ultramicroscopy* 135: 24-35

Coleman JA, Yang D, Zhao Z, Wen P-C, Yoshioka C, Tajkhorshid E, Gouaux E (2019) Serotonin transporter–ibogaine complexes illuminate mechanisms of inhibition and transport. *Nature* 569: 141-145

Crooks GE, Hon G, Chandonia J-M, Brenner SE (2004) WebLogo: A Sequence Logo Generator. *Genome Research* 14: 1188-1190

Gouaux DYaE (2021) Illumination of serotonin transporter mechanism and role of the allosteric site. *Sci A*

Harini Krishnamurthy1 & Eric Gouaux1, Krishnamurthy H, Gouaux E (2012) X-ray structures of LeuT in substrate-free outward-open and apo inward-open states. *Nature* 481: 469-74

Kawate T, Gouaux E (2006) Fluorescence-detection size-exclusion chromatography for precrystallization screening of integral membrane proteins. *Structure* 14: 673-81

Li Y, Wang X, Meng Y, Hu T, Zhao J, Li R, Bai Q, Yuan P, Han J, Hao K, Wei Y, Qiu Y, Li N, Zhao Y (2024) Dopamine reuptake and inhibitory mechanisms in human dopamine transporter. *Nature*

Singh SK, Yamashita A, Gouaux E (2007) Antidepressant binding site in a bacterial homologue of neurotransmitter transporters. *Nature* 448: 952-6

Wang KH, Penmatsa A, Gouaux E (2015) Neurotransmitter and psychostimulant recognition by the dopamine transporter. *Nature* 521: 322-7

Wei Y, Li R, Meng Y, Hu T, Zhao J, Gao Y, Bai Q, Li N, Zhao Y (2024) Transport mechanism and pharmacology of the human GlyT1. *Cell* 187: 1719-1732 e14

Yamashita A, Singh SK, Kawate T, Jin Y, Gouaux E (2005) Crystal structure of a bacterial homologue of Na+/Cl--dependent neurotransmitter transporters. *Nature* 437: 215-23

Zhang H, Yin Y-L, Dai A, Zhang T, Zhang C, Wu C, Hu W, He X, Pan B, Jin S, Yuan Q, Wang M-W, Yang D, Xu HE, Jiang Y (2024) Dimerization and antidepressant recognition at noradrenaline transporter. *Nature*

Zhu A, Huang J, Kong F, Tan J, Lei J, Yuan Y, Yan C (2023) Molecular basis for substrate recognition and transport of human GABA transporter GAT1. *Nature Structural & Molecular Biology* 30: 1012-1022
